# PEARL: Protein Eluting Alginate with Recombinant Lactobacilli

**DOI:** 10.1101/2024.09.12.612671

**Authors:** Varun Sai Tadimarri, Marc Blanch-Asensio, Ketaki Deshpande, Jonas Baumann, Carole Baumann, Rolf Müller, Sara Trujillo, Shrikrishnan Sankaran

## Abstract

Engineered living materials (ELMs) made of bacteria in hydrogels have shown considerable promise for therapeutic applications through controlled and sustained release of complex biopharmaceuticals at low costs and with reduced wastage. While most therapeutic ELMs use *E. coli* due to its large genetic toolbox, most live biotherapeutic bacteria in development are lactic acid bacteria due to native health benefits they offer. Among these, lactobacilli form the largest family of probiotics with therapeutic potential in almost all sites of the body with a microbiome. A major factor limiting the use of lactobacilli in ELMs is their limited genetic toolbox.

In this study, we build upon our recent work to expand the genetic programmability of probiotic *Lactiplantibacillus plantarum* WCFS1 for protein secretion and encapsulate it in a simple, cost-effective, and biocompatible core-shell alginate bead to develop an ELM. We demonstrate the controlled release of recombinant proteins, even up to 14 days from this ELM, thereby terming it PEARL - Protein Eluting Alginate with Recombinant Lactobacilli. Notably, lactobacillus encapsulation offered benefits like bacterial containment, protein release profile stabilization, and metabolite-induced cytotoxicity prevention. These findings demonstrate the mutual benefits of combining recombinant lactobacilli with alginate for the controlled and sustained release of proteins.

## INTRODUCTION

Engineered living materials (ELMs) are composites made of living entities integrated with materials. [1,2] The living entities in ELMs enable them to be designed with life-like abilities of adaptation, regeneration, biosynthesis, growth, etc. Additionally, the possibility to genetically reprogram these abilities to optimally suit a desired application has propelled the rapid development of ELMs in various domains ranging from architecture to medicine. ELMs have advanced in the medical field, especially due to the benefits they offer to deliver therapeutics. [3,4,5] Since the human body naturally supports microbial survival, ELM activities can potentially be sustained in the long term. The microbes can be engineered to produce high-value biopharmaceutical drugs (proteins, peptides, biomacromolecules) for delivery at the disease site. In most studies conducted to date, *E. coli* has been the predominant organism encapsulated within ELMs. For example, an ELM, termed PATCH, encapsulated lysostaphin-secreting *E. coli* to inhibit the growth of *Staphylococcus aureus*, including a methicillin-resistant *S. aureus* (MRSA) strain. [6] In another study, an *E. coli* strain engineered to light-responsively produce an anti-bacterial compound, deoxyviolacein. After encapsulation, the strain demonstrated drug release over 40 days. [7] This approach avoids the need to externally produce, purify, encapsulate, and store these often-fragile biopharmaceuticals, thereby reducing the costs and waste associated with these processes.

To achieve these benefits, therapeutic ELMs primarily use bacteria as their living entity, since they grow fast, have simple nutrient requirements, can be encapsulated at high densities, and are often naturally beneficial to the host in the form of probiotics. In fact, bacteria have been developed for over two decades as engineered live biotherapeutics that can be administered directly within the body to produce and deliver drugs on-site. [8] While clinical trials have shown that the engineered bacteria are safe for use in humans, proving their efficacy has been a major hurdle blocking their approval as therapeutic products. [9] A fundamental challenge in achieving therapeutic efficacy is ensuring that the right density of the engineered bacteria is viable and active when they reach the target site. Furthermore, this must be balanced with the need for biocontainment of the bacteria, preventing them from invading unwanted sites in the body or environmental niches where they can potentially disturb the local ecology. ELMs satisfy these needs since they can provide a conducive and protected environment for the bacteria to grow to large population sizes while physically confining them within the material. [10,11,12,13] Most therapeutic ELMs are based on probiotic *E. coli* and only a few studies cover other probiotic or GRAS (generally recognized as safe) strains like *Lactococcus lactis*, [14] *Bacillus subtilis* [15] and *Corynebacterium glutamicum.* [16] Interestingly, most engineered live biotherapeutic candidates currently in clinical trials are lactic acid bacteria. This is because lactic acid bacteria are the largest group of commensals and probiotics, beneficial not only for the gut but also other organs hosting a microbiome. In particular, the *Lactobacillus* family has the largest number of probiotic strains, many of which are being explored for therapeutic applications. [17] Among these, *Lactiplantibacillus plantarum* has been well investigated and is the most studied strain till date.[18] Several *L. plantarum* strains have GRAS status and none are reported as pathogenic to date, whereas only *E. coli* Nissle 1917 is probiotic with GRAS status and almost all other strains are pathogenic or endotoxic. Because of this, beneficial effects of *L. plantarum* strains have been demonstrated in multiple sites of the body beyond the gut, like the oral cavity, female reproductive organs, bladder, skin, etc., whereas the applicability of *E. coli* is largely limited to the gastrointestinal tract. Even there, *L. plantarum*’s thick cell wall and metabolic robustness enable it to survive harsh conditions like low pH, lytic enzymes, and bile salts, better than *E. coli*. Overall *L. plantarum* for ELMs offer significant advantages in terms of safety, health benefits, biocompatibility, and robustness. However, ELMs containing lactobacilli have not yet been reported despite these benefits, possibly due to their limited genetic programmability.

Recently, a few research groups, including ours, have made major strides in expanding the genetic programmability of *L. plantarum* WCFS1. [18] We developed a direct cloning methodology to speed up the plasmid-based engineering of this strain by circumventing the need for an intermediate host. [19] Using this strategy, we have identified the strongest constitutive promoter and repressor reported to date and characterized several toxin-antitoxin systems for antibiotic-free plasmid retention. [20,21] Other groups have reported using sakacin-inducible, xylose-inducible, and lactose-inducible systems to express heterologous proteins in *L. plantarum* WCFS1. [22] To facilitate the secretion of these proteins, various endogenous and exogenous signal peptides have been explored. [23] In addition to secretion, the anchoring of these secreted proteins has garnered significant interest in recent years, particularly in the development of bacterial vaccines. In this context, antigens have been secreted and anchored to various components of the bacterial cell surface using endogenous LysM and LPXTG moieties, enhancing their potential as vaccine candidates. [24] Nevertheless, the performance and containment of these engineered bacteria within biocompatible materials under physiologically relevant conditions is yet to be shown. Encapsulation of *L. plantarum* in hydrogels and food-based materials has been done for their delivery as probiotics. [25,26] These materials were not designed to contain the bacteria but rather to deliver them in the gut.

In this study, we have investigated how *L. plantarum* WCFS1 can be securely contained within a hydrogel. To do this, core-shell alginate beads were fabricated using a simple manual strategy that can be easily adapted by other labs even without experience in hydrogel fabrication. Interestingly, we observed that alginate in this format is particularly well suited to contain *L. plantarum* while maintaining its viability for at least 14 days, while *E. coli* was found to leak out within a day. We further expanded the genetic programmability of this bacterium by optimizing genetic parts needed to drive high-level secretion of a nuclease protein. This optimization involved identifying and testing multiple signal peptides to determine which ones most compatibly function with our recently discovered strong constitutive promoters. Encapsulation of the engineered lactobacilli in alginate beads helped to stabilize the release profiles of the secreted protein and sustained this activity over at least 14 days. Due to this prolonged protein elution capability, we hereafter term this ELM as PEARL - Protein Eluting Alginate with Recombinant Lactobacilli. With these PEARLs, we also demonstrate the secretion of two therapeutically relevant proteins, α-Melanocyte-stimulating hormone (α-MSH) and elafin, a neutrophil elastase inhibitor. Finally, encapsulation additionally aids in preventing cytotoxicity that can potentially be caused by metabolites of *L. plantarum* as confirmed by biocompatibility tests performed using mammalian cells and highly sensitive Zebrafish embryos. These results set the stage for the development of PEARLs towards therapeutic applications.

## EXPERIMENTAL SECTION

### Strain, media and plasmids

*L*. *plantarum* WCFS1 strain was used in this study. De Man, Rogosa and Sharpe (MRS) media (Carl Roth GmbH, Germany, Art. No. X924.1) was used to grow the strain. Engineered *L*. *plantarum* WCFS1 strains were grown in MRS media with 10 μg/mL of erythromycin (Carl Roth GmbH, Art. No. 4166.2) at 37°C and 250 revolutions per minute (rpm) for 16 to 20 h. *E. coli* Nissle 1917 was maintained in Luria-Bertani (LB) media (Carl Roth GmbH, Art. No. X968.1). Engineered *E*. *coli* Nissle 1917 was grown in LB media supplemented with 200 μg/mL of erythromycin at 37°C, 250 rpm for 16 to 20h. The pLp_3050sNuc plasmid was the backbone vector used in this study (Addgene plasmid # 122030).

### Molecular cloning reagents

The following enzymes and kits were purchased from NEB: Q5 High Fidelity 2X Master Mix for PCRs, NEBuilder® HiFi DNA Assembly Cloning Kit for Gibson Assembly, Quick Blunting Kit for DNA phosphorylation and the T4 DNA Ligase enzyme for DNA ligation. DNA oligos were purchased from Eurofins Genomics GmbH (Köln, Germany) (**Table S1**). Synthetic genes were synthesized as eBlocks from Integrated DNA Technologies (IDT) (Coralville, USA) and were codon optimized using the IDT Codon Optimization Tool. Plasmids were extracted using the plasmid extraction kit from Qiagen GmbH (Hilden, Germany). The DNA purification kit was purchased from Promega GmbH (Walldorf, Germany). The ladder used for agarose gels was the Generuler 1 Kb DNA Ladder from Thermo Fisher Scientific.

### Plasmid construction

We performed the direct cloning method for *L. plantarum* WCFS1, previously described by us, to assemble and obtain enough plasmid amount required for a successful transformation. [19] In brief, we used Gibson Assembly to assemble the synthetic genes into the plasmid backbone. Instead of transforming the assembled product into an E*. coli* cloning strain, we performed a PCR-based plasmid amplification, followed by phosphorylation and ligations of the PCR products for circularization. Finally, we performed a final purification to concentrate the circular DNA before *L. plantarum* WCFS1 transformation. For sequence verification, the gene of interest was amplified by PCR (Q5 high-fidelity polymerase), and the product was sent for sequencing at Eurofins Genomics GmbH (Köln, Germany).

### *L. plantarum* WCFS1 transformation

*L. plantarum* WCFS1 electrocompetent cells were prepared as described in [19]. *L. plantarum* WCFS1 transformation was done following the protocol described [19].

### Media selection

When inoculated from glycerol stock, *L. plantarum* was always grown in MRS media. To assess the containment of bacteria within PEARLs, both MRS and LB media were used for subculturing and incubation. For all NucA detection studies, bacteria were then subcultured in DNase media. For elafin and α-MSH experiments, DMEM media was used for subculturing of bacteria and incubation of PEARLs. For fibroblast experiments, fibroblast growth media was used for subculturing of bacteria and incubation of PEARLs. For zebrafish experiments, 0.3x Danieu’s media was used for subculturing of bacteria and incubation of PEARLs.

### NucA Assay (in DNase media with Methyl green)

#### Qualitative Assay

To analyze the secretion of NucA by genetically modified bacteria, the bacteria were first grown overnight in MRS media supplemented with 10 µg/mL erythromycin at 37°C, 250 rpm and the OD_600_ was measured the following day. The samples were subcultured in 4 mL of antibiotic-supplemented fresh MRS media at an initial OD_600_ of 0.1 and incubated at 37°C until an OD of 1. Then, 10 µL of the undiluted bacterial cultures were spotted on DNase agar (Altmann Analytik GmbH, Germany) supplemented with 10 µg/mL of erythromycin and incubated at 37°C for 24 h. The discoloration zone surrounding the spotted area, indicative of NucA activity was captured and documented using the BioRad Gel Documentation System.

#### Quantitative Assay

To measure the concentration of the secreted NucA, overnight bacterial cultures (in MRS broth) were subcultured to an OD_600_ of 0.1 in MRS broth. Once the OD reached 1, bacteria equivalent to OD 0.1 were transferred to an autoclaved DNase media supplemented with erythromycin of 1 mL volume and incubated at 37°C and 250 rpm shaking conditions for 24 h. The DNase media was obtained by agar precipitation of the commercial DNase Agar and the final formulation comprised Tryptose (20 g/L), Sodium Chloride (5 g/L), Calf Thymus DNA (2 g/L), Methyl Green (0.05 g/L). After 24 h, the bacterial cultures were centrifuged at 13000 rpm (15700 × g) and 200 µL of the respective supernatants were added to the clear bottom 96-well microtiter plate (Corning® 96 well clear bottom black plate, USA). The samples were analyzed in the Microplate Reader Infinite 200 Pro for methyl green-DNA complex-based fluorescent intensity (Exλ / Emλ = 633/668 nm). The z-position and gain settings for recording the fluorescence were set to 19000 µm and 100, respectively. The raw data obtained was correlated and calculated based on the standard curve explained below.

#### Standard curve

Overnight grown culture of WCFS1 - Empty vector (EV, bacteria carrying a plasmid based on the p256 origin of replication and the erythromycin resistance). was diluted to the OD 0.1 in DNase agar + methyl green media. Using the 0.1 OD DNase media prepared in the previous step as the solvent, serial dilutions of NucA protein (Sigma Aldrich) were made. In our experiment, the standard curve was plotted in the range of 200 nM - 1.56 nM (200, 100, 50, 25, 12.5, 6.25, 3.12, 1.56 nM). The final volume of each concentration was 1 mL and was prepared in 1.5 mL Eppendorf tube. Once the protein dilutions were transferred to the Eppendorf tube, they were covered with aluminum foil to protect the bleaching of methyl green from exposure to light. The tubes were stacked in a 37°C incubator with 250 rpm shaking for 24 hours.

After 24 hours, 200 µL from each Eppendorf/concentration were pipetted in duplicates into a black 96-well plate with transparent bottom. The fluorescence of samples was measured at Ex/Em of 633/668 nm at Z position of 19000 µm and Gain of 100 in the Microplate Reader Infinite 200 Pro (Tecan Deutschland GmbH, Germany).

### Flow cytometry

The fluorescence intensity of the reporter proteins mCherry and mScarlet3 in bacterial cultures was measured using the Guava easyCyte BG flow-cytometer (Luminex, USA). Samples were prepared as described in [20]. The Luminex GuavaSoft 4.0 software for EasyCyte was employed for the analysis and of the data.

### PEARL fabrication - Encapsulation in alginate core-shell hydrogels

#### Fabrication of alginate cores

3 wt% alginate solution was prepared by dissolving sodium alginate (Sigma-Aldrich 71238) in sterile MilliQ water followed by sterilization by autoclaving it at 120°C for 15 minutes. A bacterial culture (approximately 10^8^ cells in LB media) was mixed with 3 wt% alginate solution in a 1:2 volume ratio to obtain a bacterial-alginate core solution of 2 wt%. The bacterial culture was supplemented with 1% v/v Alcian blue to enhance the visualization of alginate-bacteria core beads during manual fabrication. The bacteria-alginate core mixture was vortexed thoroughly to ensure uniformity. Droplets, each with a volume of 10 µL, were pipetted and dropped into a 5 wt% CaCl_2_ solution (Ionic crosslinker). The droplets were left to polymerize and solidify in CaCl_2_ solution for 15 minutes. Solidified core beads (roughly 2 mm in diameter) were collected and washed thrice with sterile Milli-Q water to remove the remnant CaCl_2_ solution and bacteria on the surface of the beads.

#### Shell coating

1.5 wt% alginate solution for shell coating was prepared by mixing 3 wt% alginate stock solution and Milli-Q water in a 1:1 volume ratio followed by vortexing. Alginate cores from the last step of the previous section were dipped into the shell solution. Alginate bead surrounded by shell solution (roughly 25 µL) was collected with a pipette tip, whose tip was cut with a sterile blade to increase the size of the opening, followed by dropping into the cross-linker solution. The core-shell alginate beads were left to solidify for 15 minutes. The beads were collected, washed with sterile Milli-Q water, and transferred to a well plate with a nutrient media.

### Tracking changes in PEARLs using microplate reader

PEARLs (alginate beads) after fabrication were immediately transferred to a 96-well plate containing wells filled with 200 µL of respective media (LB media for OD_600_ measurements and DNase media for NucA quantification). Growth kinetics along with tracking of fluorescence expression was done for 72 hours using Tecan microplate reader with settings for Excitation/Emission wavelengths [Ex/Em for mCherry (587/625 nm) / mScarlet3 (569/600 nm) / NucA (633/668 nm), Manual gain 80-100 (depending on the magnitude of signal, a particular gain value has been selected for individual assays, between 80 and 100), Z-position 19000 µm and integration time of 20 µs]. Data was normalized and analyzed using GraphPad prism.

### Western Blot

Sodium dodecyl sulfate-polyacrylamide electrophoresis (SDS-PAGE) of denatured protein samples (Culture supernatants in our experiments) were performed. PVDF membrane (Immobilon-P, Merck Millipore) was cut to the same size as polyacrylamide gel was immersed in 99% methanol for 10 seconds followed by soaking the membrane in sterile MQ water for 10 seconds and finally dispersed in 50 mL of cold 1x Transfer buffer (25 mM Tris Base-pH 8.3, 192 mM Glycine, 200 mL methanol and rest of MQ water upto 1L) for a few minutes. Parallelly, components needed for cassette assembly (foam pad, filter paper, polyacrylamide gel) were equilibrated in cold 1x Transfer buffer for 10 minutes. The cassette was assembled in the order of Support grid/Cathode ( - ) –> foam pad –> filter paper –> polyacrylamide gel –> PVDF membrane –> filter paper –> foam pad –> Support grid/anode (+).The cassette was placed in a Bio-Rad buffer tank set on a magnetic stirrer, filled with 1x Transfer buffer, a magnetic stirring bar was placed underneath the cassette and the whole setup was connected to a power unit with the voltage set to 100 V for a duration of 90 minutes and the transfer was initiated by switching on the power supply. After 90 minutes, the PVDF membrane was peeled out (make sure that protein ladder got transferred from PAGE gel onto the PVDF membrane, indicating the successful transfer of proteins) and placed in a 50 mL amber-colored falcon by rolling the membrane inwards such that the side of the membrane with transferred proteins was in contact with the blocking buffer (15 mL of 3% Bovine Serum Albumin in 1x Tris-buffered saline (TBS)) inside. The falcon was placed in a rotary shaker and incubated at room temperature for one hour. The spent blocking buffer was discarded and 10 mL of fluorophore-tagged antibody (elafin antibody H-2 conjugated to Alexa Fluor 647 from Santa Cruz Biotechnology, in ratio of 1:500) in 3% Bovine Serum Albumin made with 1x TBS was added to the falcon. The falcon was set in a rotary shaker and incubated overnight at 4 °C. Next morning, the membrane was taken out of the falcon and washed thrice with 1x TBST (TBS supplemented with 0.05% Tween-20) followed by capturing the blot in optimized RGB settings in a Bio-Rad gel doc device.

### Neutrophil Elastase Inhibition Assay

Confirmation of elafin’s bioactivity was performed by measuring the inhibition of Neutrophil elastase (NE) enzyme. An in-house assay was optimized using human neutrophil elastase (324681-Sigma-Aldrich), neutrophil elastase specific fluorogenic substrate (MeOSuc-AAPV-AFC, from Abcam) and commercial recombinant human elafin (R&D systems). In the absence of NE inhibitors, NE recognizes & binds to specific tetrapeptide sequence (AAPV) and cleaves the bond between valine and AFC (7-amino-4-trifluoromethylcoumarin), a fluorescent moiety at the C-terminus of substrate. Upon cleavage, AFC is released and exhibits strong fluorescence which can be measured using a plate reader. In presence of elafin or any NE inhibitor, NE is reversibly or irreversibly blocked from interacting with the cleavage site of substrate and hence AFC will remain intact and its fluorescence is quenched resulting in low values.

#### Standard curve

Desired concentrations of elafin (80 nM, 40 nM, 20 nM, 10 nM, 5 nM, 2.5 nM, 1.25 nM, 0.62 nM) were prepared by serial dilution of the stock protein in DMEM media. Parallelly, 1 mU Human Neutrophil elastase (Sigma Aldrich, Germany) solution was prepared by diluting in assay buffer (200 mM NaCl, 200 mM Tris-HCl, pH 7.5, 0.01% BSA, 0.05% Tween-20) (REF). 25 µL containing 1 mU of Neutrophil elastase was pipetted into a Black 96-well plate with a flat transparent bottom (Greiner Bio-one, 655096). 50 µL of serially diluted elafin solution with a specific concentration was added to wells with neutrophil elastase and pipetted to ensure the mixing of both components. The plate was transferred to a 37 °C static incubator for 10 minutes and taken out. 25 µL of 100 µM fluorogenic elastase substrate (MeOSuc-AAPV-AFC, from Abcam) was added to the wells to make a final volume of 100 µL. The plate was immediately transferred to a plate reader and the fluorescence of the samples was measured at excitation/emission of 380/460 nm (Gain = 70, Z-position = 17000 µm).

#### Detection from culture and PEARLs

50 µL of culture supernatant was added to the wells with 25 µL of neutrophil elastase (NE) (1mU of NE), pipetted to ensure mixing, and incubated at 37 °C for ten minutes. 25 µL of elastase substrate (of 100 µM concentration) was added to make a final volume of 100 µL. The plate was immediately transferred to a plate reader (Microplate reader Infinite 200 Pro) and the fluorescence of the samples was measured at excitation/emission of 380/460 nm (Gain = 70, Z-position = 17000 µm).

### Nano Luciferase Assay

To measure the amount of fusion protein with the SmBiT tag getting secreted, 25 µL of culture or PEARL supernatant was transferred to a white, flat-bottomed 96-well plate and 25 µL of LgBiT of 1µM concentration, prepared in PBS was added. The furimazine substrate (Promega, Walldorf, Germany) was diluted 1:50 in Nano-Glo Luciferase assay buffer (Promega, Walldorf, Germany) and immediately added to wells with SmBiT-LgBiT to make a final volume of 100 µL. The well plate was protected from light and incubated at room temperature for 10 minutes. Using the Microplate reader Infinite 200 Pro (Tecan Deutschland GmbH, Germany), luminescence values were measured with an integration time of 100 ms. Raw data was correlated with the standard curve obtained using synthesized SmBiT peptide and purified LgBiT fragment (**Figure S8a**) to calculate the concentration of secreted proteins.

### Detection of bacterial escape

Tracking of escape or leakage of bacteria from alginate beads was done in two ways: 1) Transfer of alginate bead onto nutrient agar plate (MRS erythromycin agar and LB kanamycin agar) immediately after fabrication, incubate in a 37 °C static incubator for up to 2 days and observe whether colony outgrowth is observed or not; 2) Take OD_600_ measurement of alginate beads immersed in 200 µL of nutrient media in a well of a 96-well plate to detect even a slight increase in turbidity of surrounding media due to bacterial escape and subsequent growth. The final values are the mean of 9 different points measured within a single well.

### Bacterial recovery from PEARLs

To determine the viable population of bacteria within the alginate core after 14 days, alginate macro beads were treated with 200 µL 25 mM of EDTA (pH 8) in a 1.5 mL Eppendorf for 30 minutes with shaking in a thermomixer (30 °C, 1000 rpm). The mixture containing EDTA, dissolved alginate, and bacteria was centrifuged at 8000 rpm for 5 minutes. EDTA supernatant was discarded, bacterial pellet was resuspended in 100 µL of nutrient broth and plated on an agar plate supplemented with erythromycin (10 µg/mL). The agar plate was incubated in a 37 °C static incubator for 48 hours followed by imaging and automated counting of colonies to determine CFU/mL.[27] Automated counting involved adjusting the threshold of 8-bit images of agar plates such that only bacterial colonies were selected excluding outlines and edges, followed by the selection of analyze particles function resulting in the pop up window with results including location and number of colonies in agar plate) of grown bacterial colonies in an agar plate was done after ours of incubation (at 37 °C) using ImageJ.

### Evaluation of the cytocompatibility of PEARLs

#### Cell culture

Normal human dermal fibroblast (NHDF) cells were purchased from Lonza, Germany. The cells (passages between 10-12) were cultured in growth media (FBM™ Fibroblast Growth Basal Media (Lonza, Germany) supplemented with 1% penicillin−streptomycin (P/S) and FGM-2 SingleQuots supplements (containing insulin, hFGF-B, GA-1000, 2% FBS)). All cells were incubated at 37°C and 5% CO_2_. The media was exchanged every other day. The cells were sub-cultured after growing to 60−80% confluency.

#### PEARL supernatants for cytotoxicity assays

The cytocompatibility of PEARLs was evaluated in two ways. First using supernatants (200 μL in FGM) from PEARLs following their incubation in FGM in a 37 °C static incubator were collected after 24 hours and used to treat the NHDFs and then, performing a co-culture of the PEARLs together with a monolayer of NHDFs. Cells (34,000 cells/well) were seeded in sterile µ-Plate Angiogenesis 96 well-plates (Ibidi, Germany) in growth media (70 μL) and allowed to attach overnight (16 h, 37°C, 5% CO_2_). Then, the media was removed, cells were washed twice with DPBS and either treated with supernatants from the PEARLs or treated with the PEARLs for co-culture. Conditions were incubated overnight at 37°C, 5% CO_2_. The conditions investigated were: “Alginate” (supernatants from empty gels without bacteria), “WT” (supernatants from wild-type bacterial gels), “*P_tec_*_3050” (supernatants from high-secreting NucA bacterial gels), “*P_tlpA_*_3050” (supernatants from low secreting NucA bacterial gels), “Alginate+Nuc” (supernatants from alginate gels loaded with 50 nM of NucA), “Nuc control” (media containing 0.5 nM, 5 nM or 50 nM of NucA).

#### Cytotoxicity assays

Lactate dehydrogenase (LDH) assay (CytoTox 96 Non-Radioactive Cytotoxicity Assay, Promega) was performed to quantify the percentage of cell death after treatment following the manufacturer’s instructions. Briefly, 30 μL of supernatant from each well was transferred to a new 96-well plate, and 30 μL of LDH substrate was added. Samples were incubated in the dark for 30 min at room temperature. Then, 30 µL of the stop solution was added and the absorbance at 490 nm was read using a plate reader (TECAN Spark). A negative control of cells cultured on cell growth media, and a positive control of cells treated with 37.5% Triton X-100 (lysis control) were also analyzed. The blanks used were media background absorbance readouts at 490 nm. The percentage of cell death was calculated as:

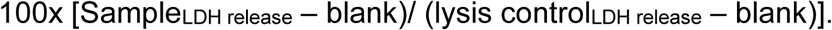

#### Cell viability assays

AlamarBlue assay (Invitrogen) was used to measure cell viability after treatment, following the manufacturer’s guidelines. Briefly, alamarBlue reagent (10% v/v in growth media) was added to treated cells (cells incubated for 24 h with bacterial alginate beads in co-culture or their supernatants) and incubated for 2 h. The media was then transferred to black 96-well plates. Fluorescence (Ex/Em 570/600 nm) was analyzed using a plate reader (TECAN Spark). Controls of cells cultured on cell growth media and blanks with only media were also analyzed. To calculate cell viability, fluorescence values were normalized with the values of the control as follows:

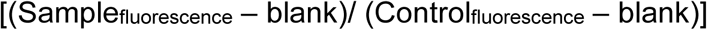

#### Cell staining

Cells were fixed with 4% paraformaldehyde (PFA, in DPBS) for 20 minutes at room temperature and washed twice with DPBS. Then, cells were permeabilized with 0.1% Triton X-100 (in DPBS) for 5 minutes. After permeabilization, samples were washed twice with DPBS. Then cells were blocked using blocking buffer (1% bovine serum albumin (BSA) in DPBS). For actin staining, cells were incubated with alexaFluor 488-Phalloidin (dilution 1:400 in blocking buffer), and for nuclei staining, cells were stained with 4′,6-diamidino-2-phenylindole (DAPI) (dilution 1:1000 in blocking buffer) for 1h at room temperature and protected from light. After staining, samples were washed three times with DPBS. Images were taken using a Leica DMI6000 B epifluorescence microscope at 20X magnification. Image analysis was carried out using ImageJ software (version 1.53k). Nuclei were counted using ImageJ to quantify cell density from the images. Images were converted to 8-bit and the Gaussian blur filter was selected with a ball radius of 2. After that, the find maxima tool was applied with a noise tolerance of 20. Maxima points found were counted (one per nucleus detected in the image). Finally, the number of nuclei per area was calculated.

### Zebrafish embryo-based Assays

#### Zebrafish Strain and Maintenance

Husbandry of adult zebrafish (ZF) was carried out in accordance with internal guidelines based on the German Animal Welfare Act (§11 Abs. 1 TierSchG). Embryos of wild-type ZF from the AB strain used for all investigations originated from the HIPS in-house zebrafish facility. The AB strain was originally purchased from the European Zebrafish Resource Center at Karlsruhe Institute of Technology (EZRC, KIT Karlsruhe, Germany). All experiments were carried out before 5 days post-fertilization (dpf) as these early devolvement stages are not considered as animal experiments according to the EU Directive 2010/63/EU. [28] ZF embryos were maintained at 28 °C in fresh 0.3× Danieau’s media (17.4 mM NaCl, 0.21 mM KCl, 0.12 mM MgSO_4_, 0.18 mM Ca(NO_3_)_2_, 1.5 mM HEPES, 1.2 µM methylene blue, pH 7.1–7.3) until they were used for the tests. At a maximum of 5 dpf, all embryos were euthanized by hypothermal shock and consecutive freezing.

#### Biotests

Three distinct types of alginate beads were produced based on the encapsulated bacterial strains: 1) eight alginate beads containing *L. plantarum* wild type, 2) eight PEARLs with *L. plantarum P_tec__*3050_NucA (which secretes the highest level of NucA), 3) eight empty alginate beads without bacteria in the core and 4) eight PEARLs with *L. plantarum P_tec__*3050_NucA (which secretes the highest level of NucA) preincubated for 4 days in the ZF media prior to test start in modified 0.3x Danieau’s media (pH 8 with 1% v/v sucrose) to assess the effect of maximum secretion of bacterial secretome on ZF embryos. These beads were fabricated in the morning on the day of experiment and placed in a flat-bottom 48-well plate and incubated in modified 0.3× Danieau’s media to maintain pH stability and promote bacterial growth. As a positive control, culture supernatant from *L. plantarum* (OD_600_ 1 equivalent) was incubated in modified 0.3× Danieau’s media for 96 hours at 37°C with shaking, to evaluate the effect of the highest concentration of non-encapsulated bacterial secretions on ZF embryos. The 48-well plate, containing alginate beads immersed in 350 µL of modified 0.3× Danieau’s media per well, was further incubated till the addition of three zebrafish embryos per well for subsequent testing.

At 1 dpf ZF embryos were de-chorionated by treating the eggshell with pronase (1 mg/mL; Sigma, P5147-1G, lot: SLCF6250) for approx. 6 minutes. When the chorion started to disintegrate, the digestion process was stopped by washing the embryos with fresh Danieau’s media 3-times, which did not contain methylene blue in order to prevent inhibition of *L. plantarum* in the following test. Three de-chorionated embryos each were transferred to a well pooled in a volume of 50 µL pure media. The survival as well as sub-lethal effects such as morphological, physiological and behavioral alterations were investigated with a microscope and recorded at test start and after 24, 48, 72, and 96 hours (test end) corresponding to 1, 2, 3, 4 and 5 dpf, respectively. The following conditions were tested: a negative control containing only pure enriched Danieau’s media, empty alginate beads with no bacteria in the core, alginate beads containing wild-type *L. plantarum*, PEARLs containing *L. plantarum* harboring *P_tec__*3050_NucA and the last ones being alginate beads with *L. plantarum P_tec__*3050 NucA pre-incubated for 4 days prior to test start on the plate. Each condition was tested in 8 replicates, each containing 3 embryos resulting in 24 embryos for each condition.

In a separate test, three different concentrations of cell-free supernatant serving as positive control of bacterial secretome were tested, the pure supernatant as well as two dilutions containing 50 and 25 %, and a negative control containing only pure modified Danieau’s media. The supernatant was diluted using a modified Danieau’s media. The test was performed in 96-well plates with each well containing 300 µL of test solution and one embryo each. Each condition was tested in 20 replicates.

All tests were performed in parallel using ZF embryos from the same batch.

### Bioinformatics analysis

InterProScan was used to identify the signal peptides within proteins encoded in the genome of *L. plantarum* WCSF1. Signal P6.0 was used to predict the signal-peptide-driven secretion of NucA.

### Statistical analysis

Statistical analysis was done using GraphPad Prism 7.0 software. Student’s T-tests were used to determine significant differences between the means of the groups. For the cell experiments, the goodness of fit of the data was tested via Normality Shapiro-Wilk test. Normal distributed populations were analyzed using analysis of variance (ANOVA) test followed by a Tukey’s *post hoc* test to correct for multiple comparisons. Non-Gaussian populations were analyzed via the non-parametric Kruskal-Wallis test followed by a Dunn’s *post hoc* test to correct for multiple comparisons.

Differences among groups are indicated as: *p*-values <0.05 (**\***), *p*-values <0.01 (**\*\***), *p*-values <0.005 (**\*\*\***), *p*-values <0.001 (**\*\*\*\***), and differences among groups not statistically significant (**ns**).

## RESULTS AND DISCUSSION

### 1) PEARL fabrication and characterization of encapsulated *L. plantarum*

To study the behavior, secretory performance, and biocompatibility of encapsulated probiotic lactobacilli, we developed the PEARL format, which is an easy-to-fabricate alginate core-shell macro-bead ELM as shown in **Figure 1a**. *L. plantarum* WCFS1 was encapsulated in an alginate bead by manually dropping a viscous alginate-bacteria mix into a calcium chloride bath to form the core gel as previously described. [29] A polysaccharide-staining dye, Alcian Blue, was added to this mix to improve the visibility of the core gel for the next step. These beads were then submerged in a viscous alginate solution. The core bead surrounded by the alginate solution was then dropped into the calcium chloride bath to form a thick shell layer (**Figure 1a**).

**Figure 1:**
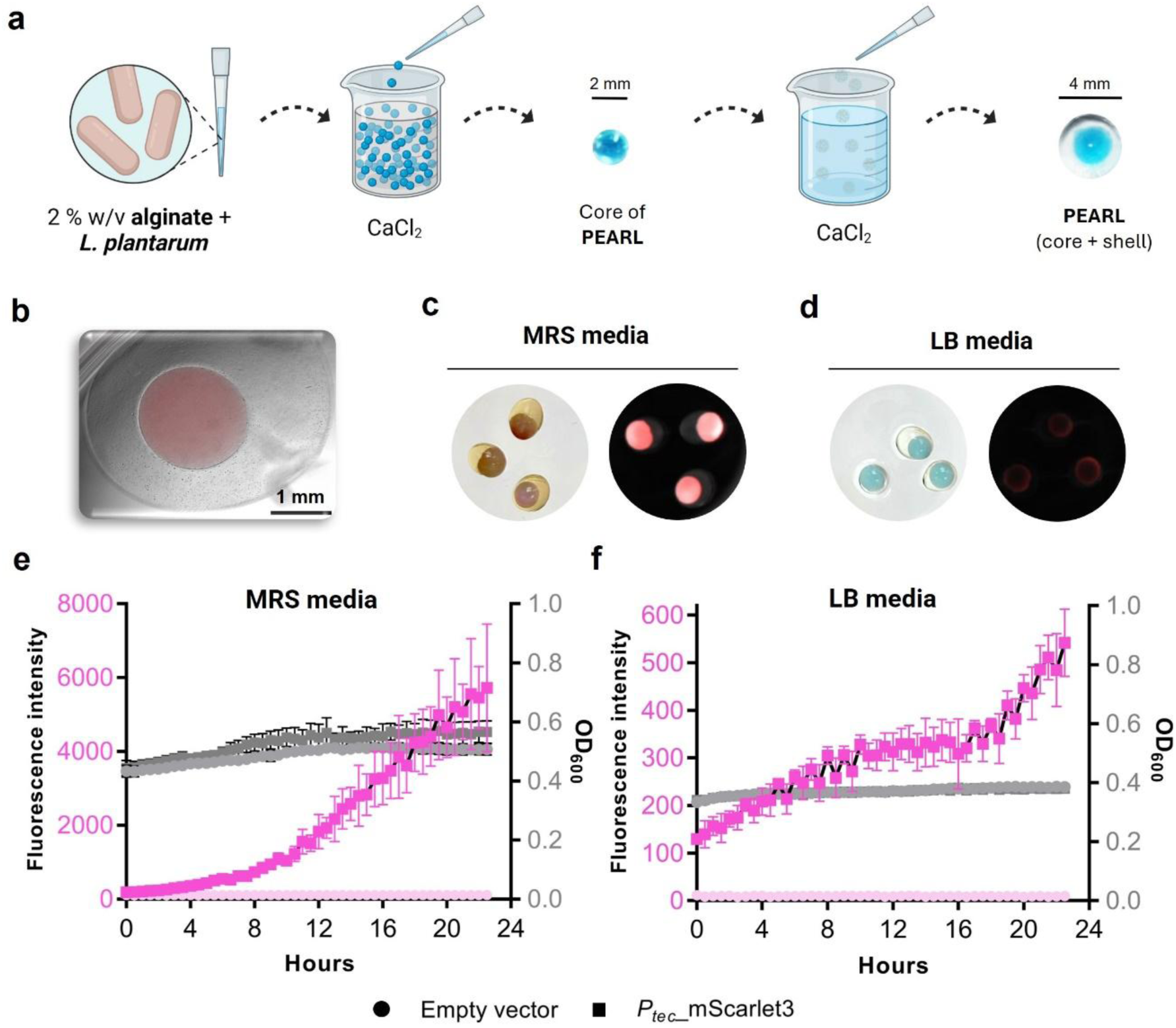
Lactobacilli-based alginate macro beads design and fabrication: **a)** Schematic representation of manual fabrication of alginate-based PEARL. It depicts the step-by-step making of PEARL. **b)** 4x overlay microscopic image of the PEARL with a core comprising mScarlet3 expressing *L. plantarum* and a shell surrounding the core. **c)** Macroscopic images of PEARLs after 24 hours incubation in MRS media; To the left is the brightfield image depicting the bacterial core and shell, and to the right is the CH3 fluorescent image showing the strong fluorescence of mScarlet3 in the core. The circular field of view is ∼ 15 mm. **d)** Macroscopic images of PEARLs after 24 hours incubation in LB media; To the left is the brightfield image depicting the bacterial core and shell, and to the right is the CH3 fluorescent image showing the fluorescence of mScarlet3 in the core. The circular field of view is ∼ 15 mm. **e)** 24-hour kinetic measurements of fluorescence intensity and OD_600_ values of encapsulated *L. plantarum* expressing mScarlet3 in PEARLs grown in MRS media (n=4, mean ± SD). **f)** 24-hour kinetic measurements of fluorescence intensity and OD_600_ values of encapsulated *L. plantarum* expressing mScarlet3 in PEARLs grown in LB media (n=4, mean ± SD).

To study the growth and containment of probiotic lactobacilli within PEARLs, *L. plantarum* WCFS1 was engineered to constitutively express mScarlet3, a highly bright and stable fluorescent protein [30]. For that, we exchanged mCherry for the mScarlet3 gene (**Figure S3a**), and we could visually confirm an increase in the pellet brightness intensity by eye (**Figure S3b**) and flow cytometry (**Figure S3c**).

After the fabrication of PEARLs with roughly 10^6^ cells (equivalent to OD_600_ of 0.01) per bead, they were incubated in MRS, the optimal media, or LB media at 37 °C. We assess the performance of the PEARLs in LB since it is a non-optimal and minimal nutrient media for these bacteria. Ideally, this ELM format is meant to perform in environments where nutrients might be scarce. Therefore, we wanted to confirm the activity of bacteria within PEARLs incubated in LB media. By fluorescence microscopy, we observed that mScarlet3 fluorescence was only detected in the core of the PEARL (**Figure 1b**) indicating the secure containment of bacteria within the core. Furthermore, we could also see differences when PEARLs were incubated in either MRS or LB media. Only in MRS the core was visually colorful due to prominent levels of mScarlet3 production (**Figure 1c**). However, when the PEARLs were incubated in LB media, mScarlet3 fluorescence could be confirmed by the BioRad Gel Documentation System (**Figure 1d**).

To quantify bacterial growth and mScarlet3 expression in PEARLs incubated in both MRS and LB media, optical density at 600 nm and fluorescence intensity were measured. A slight increase in optical density at 600 nm (OD_600_) and a significant increase fluorescence intensity (Ex/Em - 569/600 nm) were measured over 24h in a microplate reader (**Figure 1e** & **1f**), indicating that bacteria are active but stay contained within the core as outgrowth into the media was not observed.

In contrast, this ELM format was unable to contain *E. coli* Nissle 1917, neither in LB agar plates (**Figure S4a**) nor in culture (**Figure S4b**) beyond 8 h, as observed by bacterial growth around the beads and the rapid increase in OD_600_ values. This indicates that alginate is particularly suited to contain *L. plantarum* when fabricated in a core-shell format.

Apart from ensuring containment, supporting biomass growth, and enabling protein expression, the PEARLs also help prevent the pH of the media from dropping below 6, which occurs when the bacteria are cultured without encapsulation **(Figure S4c)**. While the specific causes for this surprising outcome are unknown, possible reasons could include restricted diffusion of acidic byproducts through the calcium alginate matrix, [31] or potential alterations in bacterial metabolism due to the encapsulated environment.[10]

These promising results showed that *L. plantarum* could grow and produce a heterologous protein in a contained manner within the PEARLs. For application as a therapeutic ELM, it is desirable that the bacteria secrete proteins, and these proteins diffuse out of the material. In the next sections, we engineer robust protein secretion from *L. plantarum* using strong promoters recently discovered by our group and demonstrate their functionality in PEARLs. [20]

### 2) Enhancing protein secretion in *L. plantarum*

#### Identification and selection of endogenous signal peptides (SPs)

Protein secretion in bacteria is enabled by SPs at the N-terminal of peptides that are recognized by the cellular secretion machinery, which transports the proteins across the cell membrane. While several SPs have been identified in *L. plantarum*, it was necessary to identify optimal candidates compatible with our previously discovered promoters and drive higher secretion of protein of interest compared to the reported signal peptides. For this, we screened the protein sequences encoded by more than 200 endogenous genes via InterProScan until we identified 40 SPs. As a reporter protein, we chose a nuclease (NucA) from *Staphylococcus aureus*, which can be easily detected in the media using a simple DNA degradation assay and has been extensively used to characterize secretion in gram-positive bacteria. [23, 35, 36] SignalP6.0 was used to predict NucA secretion mediated by all SPs. [37] The predictions of 5 SPs with varying secretion levels, henceforth termed SP1-5 (SP1: lp_2175, SP2: lp_2959, SP3: lp_1132, SP4: lp_0819, SP5: lp_3151) were experimentally verified. We additionally included one of the most efficient and used SPs from the literature, Lp_3050 in the testing for comparison. [24] NucA secretion was assessed using DNase agar and media assays (**Figure 2a**).

**Figure 2:**
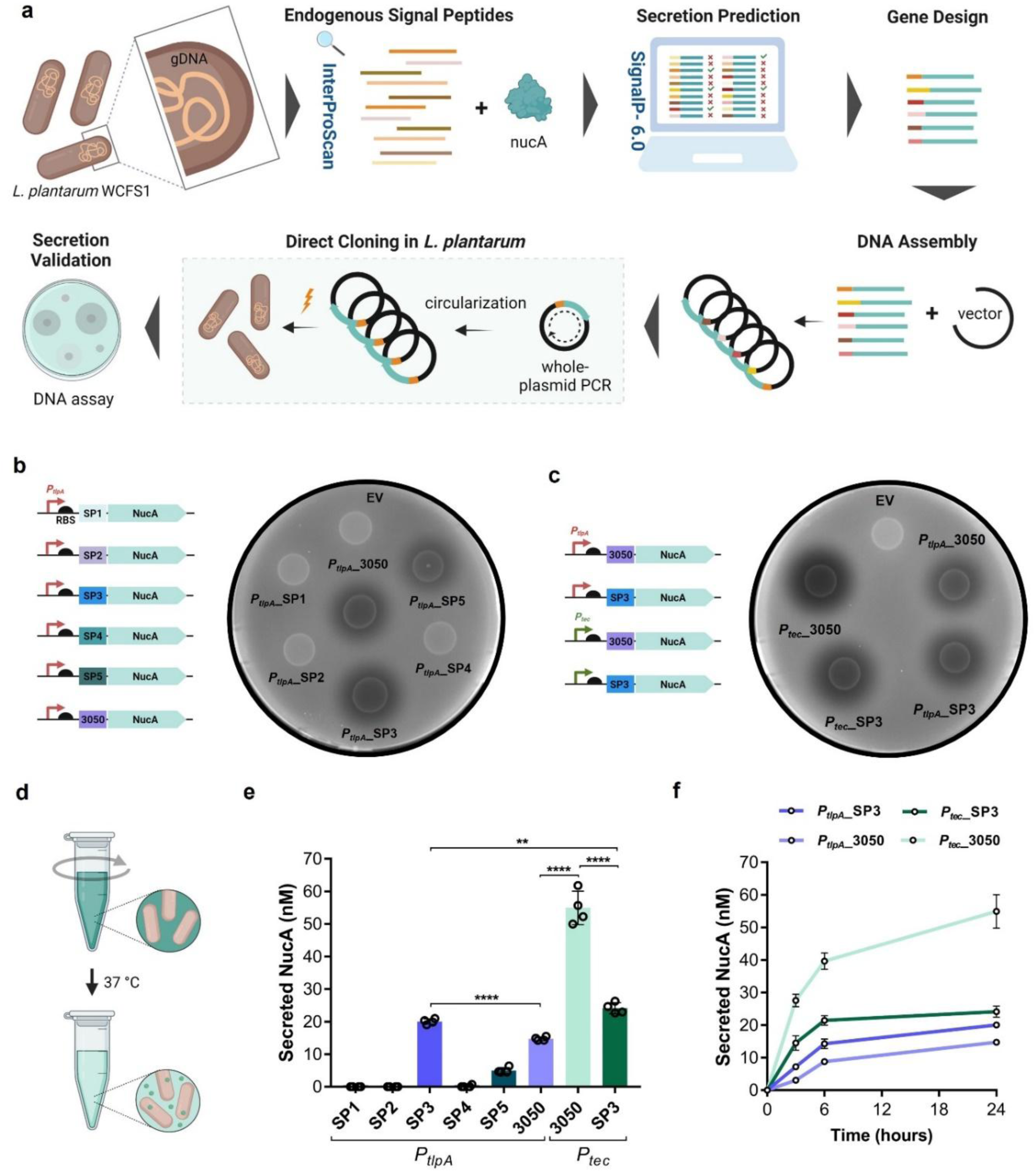
Workflow depicting quantification of recombinant protein secretion in DNase agar and DNase media: **a)** Flow scheme representing the screening of endogenous signal peptides followed by gene design and molecular cloning in *L. plantarum* WCFS1 to validate the secretion of staphylococcal NucA (17 kDa), a reporter protein commonly used for secretion studies. **b)** Schematic of genetic constructs of *P_tlpA_* promoter driving the expression of NucA, each tagged with one of six different signal peptides and EV bacteria (as negative control); to the right is the image of DNase agar plate illustrating the secretion of NucA from 10 µL of bacterial spots. **c)** Schematic of genetic constructs of *P_tlpA_* and *P_tec_* promoters driving the expression of NucA, each tagged with two best SPs (Lp_3050 and SP3) from this study; to the right is the image of DNase agar plate illustrating the secretion of NucA from 10 µL bacterial spots of these genetic constructs. **d)** Graphical representation of liquid culture-based assay to quantify the NucA secretion from engineered bacterial strains: NucA secreted from bacteria depolymerizes the DNA, thereby breaking down the methyl green-DNA complex in media resulting in loss of green coloration. **e)** Quantification of secreted NucA in DNase media after 24 hours of incubation for all the secretion-based genetic constructs screened in this study (n = 3, mean ± SD). **f)** Timepoint-based quantification (n =3, mean ± SD) (3h, 6h, 24h) of secreted NucA (in nanomolar, nM) in DNase media for two best constructs of each promoter (*P_tec_*_3050, *P_tec_*_SP3, *P_tlpA_*_3050, *P_tlpA_*_SP3).

#### Characterization of novel endogenous SPs using NucA assay

The promoters for *L. plantarum* recently discovered in our lab are *P_tlpA_* and *P_tec_* (5x and 40x stronger than previously reported). [21, 20] To avoid the risk of saturating the secretion machinery in *L. plantarum*, we initially tested the ability of the weaker promoter out of the two, *P_tlpA_*, to drive the production and secretion of NucA fused to each of the SP (SP1-5 and Lp_3050) (**Figure 2b**). Secretion was semi-quantitatively characterized using a DNase agar plate assay in which active extracellular NucA creates a dark halo around secreting colonies by degrading fluorescently labeled DNA in the agar. Only three SPs: Lp_3050, SP3 and SP5 were found to secrete detectable amounts of active NucA after 24 hours of growth on DNase agar plates. Two parameters were measured to compare the performance of each SP in the DNase agar plates - the area of the halo and the decrease in fluorescence intensity within this halo. Based on this analysis, SP3 was found to generate the biggest halo (∼4 cm^2^) and the largest drop in fluorescence (**Figure S5a**), followed by Lp_3050. Next, we tested the possibility of increasing SP3 and Lp_3050-driven secretion using the *P_tec_* promoter (**Figure 2c**). We observed that both SP3 and Lp_3050_NucA-driven secretion could be increased when *P_tec_* was the promoter driving the gene expression since bigger and darker halos (**Figure S5b**) were detected for *P_tec_* compared to *P_tlpA_*. These results proved that NucA secretion can be increased by replacing the Lp_3050 signal peptide for SP3 under *P_tlpA_* or swapping *P_tlpA_* by *P_tec_* for both Lp_3050 and SP3. Yet, the best promoter-SP combination was found to be *P_tec_*_Lp_3050.

To quantify the amount of NucA secreted by these promoter-SP combinations, we used liquid DNase media instead of the agar, in which the bacteria were allowed to grow for 24 hours at 37°C with shaking (**Figure 3d**). This enabled us to accurately measure the drop in fluorescence intensity in the media and interpolate the values in a NucA standard curve (**Figure S6a**). Once again, NucA secretion was observed only with SP3, SP5 and Lp_3050, validating the results obtained with the agar plates (**Figure 2b**). SP3 mediated better secretion than Lp_3050 when driven by *P_tlpA_* (∼20 nM and ∼15 nM, respectively). However, both SP3 and Lp_3050-driven secretion levels could significantly increase when *P_tlpA_* was swapped by *P_tec_* (∼24 nM and ∼55 nM, respectively) (**Figure 3e**). Interestingly, with *P_tec_*, Lp_3050 performed better than SP3, indicating that SPs may perform differently when expression is driven by different promoters.

**Figure 3:**
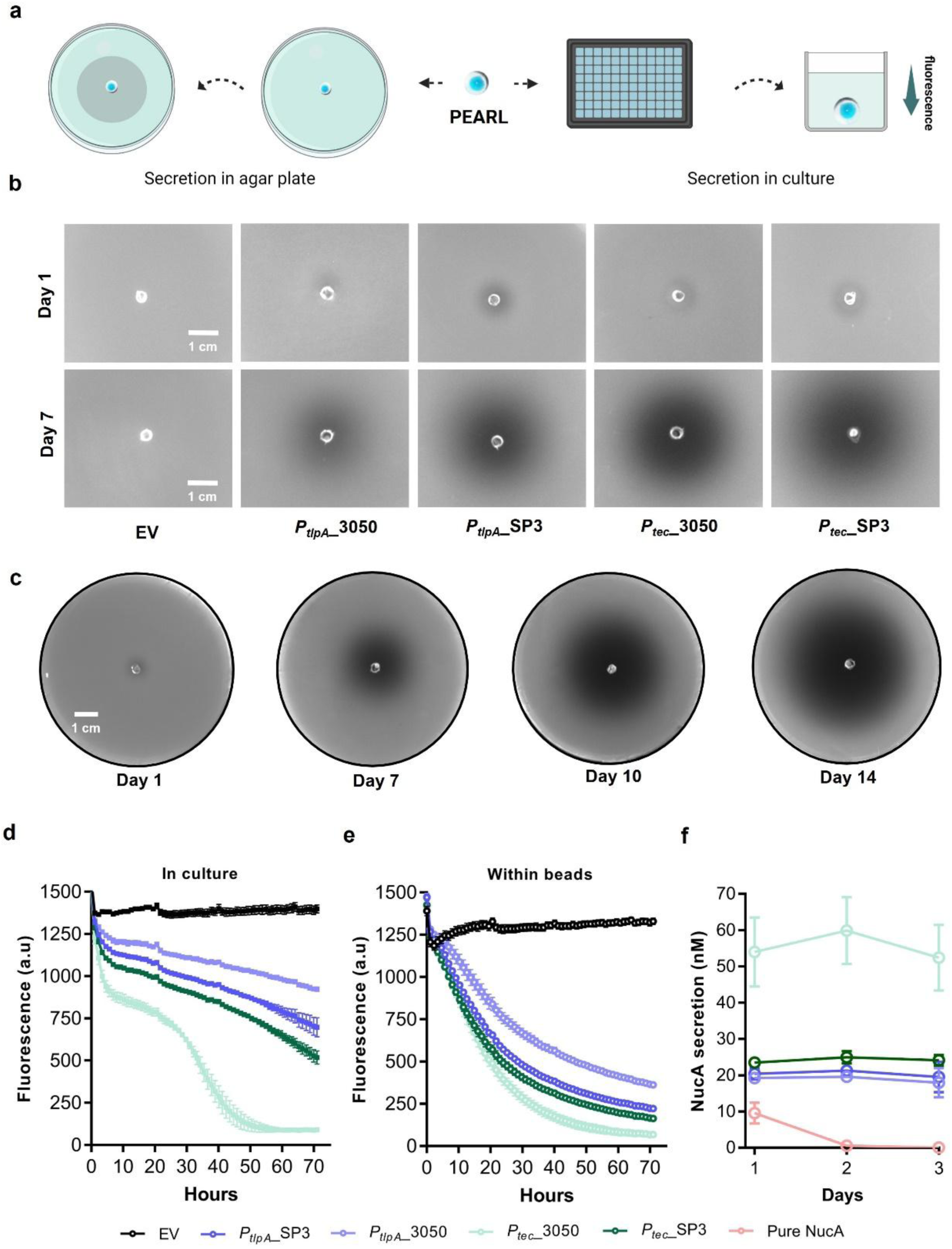
Characterization of PEARL’s release profile in DNase agar and DNase media: **a)** Graphical representation of assays done to validate and quantify the secreted NucA from alginate beads using DNase agar plate and by fluorescence kinetics in culture. **b)** DNase agar plate-based qualitative assay to track the NucA secretion and bacterial containment within PEARLs on Day 1 and Day 7. **c)** NucA secretion from PEARL embedded in DNase agar plate captured on Day 1, 7, 10, 14 indicating the bacterial viability up to 14 days **d)** Quantification of NucA secretion with reference to the drop in fluorescence in wells with culture for 72 hours (3 days) (n=5, mean ± SD). **e)** Quantification of NucA secretion with reference to the drop in fluorescence in wells with alginate beads for 72 hours (3 days) (n=5, mean ± SD). **f)** Plot showing the comparison of secreted NucA in nanomolar (n=5, mean ± SD) on Day 1, 2 & 3 from PEARLs for *P_tec_*_3050, *P_tec_*_SP3, *P_tlpA_*_3050 and *P_tlpA_*_SP3.

Based on the growth curve of *L. plantarum* WCFS1 in LB media (**Figure 1g)**, it is evident that the bacteria grow exponentially for about 6 h beyond which they reach their saturation phase due to limitation of nutrients. This change in growth phase can influence protein production and secretion, so we quantified NucA secretion at mid (3 h) and late log phase (6 h) in addition to the late saturation phase (24 h). We observed that for all the promoter/SP combinations, most of the protein (>60% approximately) is secreted within first 6 h, suggesting that the promoters are most active in the log phase of growth (**Figure 2a**). Furthermore, most of the secretion from the *P_tlpA_* constructs occurs over the first 6 h, whereas from the *P_tec_* constructs, most secretion occurs in the first 3 h (**Figure S6b**). We examined both the promoter’s activity using another reporter, mCherry, in the different growth phases, and we observed that *P_tec_* is much more active in the early and mid-log phase, which supports our results obtained with the NucA where most of the protein is produced (**Figure S6c**). Such high levels of production and secretion in the hour’s timescale are well-suited for short-term protein delivery within the body.

### 3) Assessing protein release from PEARLs

NucA release from PEARLs was assessed by two different methods: 1) by placing the PEARLs in the center of a DNase agar plate and imaging the agar plate at regular time intervals, and 2) by placing the PEARLs in a well containing DNase media followed by fluorescence measurements (**Figure 3a**). In DNase agar plates, we observed halo formation at 24 h after placing the bead in the center of the plate for all the NucA-secreting clones based on the *P_tlpA_*/*P_tec_* promoters and SP3/Lp_3050 SPs (**Figure 3b**). The halos kept increasing over the next 14 days, suggesting continuous NucA secretion from the bead. We observed the same secretion trend for the different promoter/SP combinations as observed in culture (**Figure 3b**), with the *P_tec_*_3050 strain producing the biggest and darkest halos. We also noticed that the bacteria remained contained in the bead and secretion was sustained for at least 14 days (**Figure 3c**). However, it is to be noted that since the PEARLs in this study are manually fabricated, if the core is misaligned in the shell leading to a thin shell on one edge, outgrowth of the bacteria, indicated by colonies outside the bead, is observed within 1-2 days. This occurred in approximately 10% of the PEARLs that were fabricated, and these constructs were not included in any of the analyses.

To quantify NucA secretion from the PEARLs, DNA degradation in liquid DNAse media was measured over 3 days under static incubation conditions. Non-encapsulated bacterial cultures with the same number of cells as in the alginate core of the PEARLs (10^6^) were included in the experiment for comparison. Interestingly, NucA secretion from non-encapsulated bacteria was distinctly non-monotonic, with a rapid increase within the first 8 h and then a much slower increase over the next 72 h (**Figure 3d**). In the case of *P_tec__*3050, another increase in secretion rate was observed around 24 h, and the final plateauing around 45 h is attributed to the complete degradation of the probe DNA in the DNase media. It is suspected that these different secretion profiles are related to the different growth phases that these bacteria go through during this period associated with metabolic changes and possible biofilm formation (**Figure S6c**). [38] In contrast, NucA release profiles from the PEARLs were found to be much more stable with a largely monotonic rate observed till at least 24 h followed by a gradual plateauing most likely caused by the decrease in DNA substrate availability (**Figure 3e, S7a-d**). Even the differences in protein secretion rates observed between the different promoter-signal peptide combinations seem to have considerably reduced during the first 24 h.

Furthermore, we wanted to study whether bacteria actively sustained NucA secretion for days when encapsulated in PEARLs or if most of the secretion takes place within the first day after encapsulation and the secreted NucA remains active for several days. For that, we incubated the PEARLs in DNase media, collected the media after 24 hours, measured the levels of NucA, performed several washes with PBS, and added fresh DNase media and recorded the data upto 3 days. We observed that the PEARLs secrete very similar levels of NucA day after day (**Figure 3f**). On the other hand, pure NucA encapsulated in core-shell alginate beads did not exert NucA activity outside the bead beyond the first day. This confirmed that pure NucA diffused quickly from the beads and is then washed away, whereas PEARLs keep secreting NucA (**Figure 3f**).

Considering our previous results indicating that the *P_tec_* and *P_tlpA_* are most active in the early- to mid-log phases (**Figure 2f**), the results with the PEARLs suggest that the bacteria remain in a state equivalent to these growth phases within the alginate. This is possible since bacterial growth is restricted by mechanical forces imposed by the hydrogel matrix rather than nutrient unavailability. [10, 11] Thus, while the population size is not necessarily increasing, the bacteria probably have access to nutrient levels similar to what they would have in liquid culture at the early- or mid-log phase. Thus, in the PEARL format, the most productive phase of the engineered bacteria is prolonged.

### 4) Assessing the compatibility of PEARLs for therapeutically relevant proteins

To test the compatibility of the PEARL platform to secrete therapeutically relevant bioactive proteins, we engineered *L. plantarum* to secrete elafin (also known as Trappin-2) and α-melanocyte stimulating hormone (α-MSH) as two independent secretory strains. Elafin, an 11 kDa protein, inhibits serine proteases such as neutrophil elastase and proteinase-3. By inhibiting these proteases, elafin protects tissues from protease-mediated damage in inflammatory conditions and plays a protective role in different parts of the human body, including the skin, lungs, and gastrointestinal tract. Previous studies have demonstrated the secretion of elafin from engineered *Lactococcus lactis* and *Lactobacillus casei* to explore its potential in treating inflammatory bowel disease. [32] To identify the genetic construct capable of secreting a complex protein like elafin (with 4 disulfide bonds) in *L. plantarum* WCFS1, we tested the same circuit combinations as previously tested for NucA. Using the *P_tec_* promoter, we could not obtain a mutation-free construct of elafin, likely due to the high strength of the promoter causing metabolic stress in bacteria to produce such complex protein. However, with the *P_tlpA_* promoter, the cloning was possible without mutations and we detected the secretion of elafin in combination with the SP3 but not with Lp_3050 (Western blot, **Figure S9b**). Therefore, we used the *L. plantarum* strain harboring *P_tlpA_*___SP3_elafin construct for further experiments to demonstrate the functionality of the PEARL system.

Western blot analysis of denatured culture supernatants demonstrated successful secretion of elafin in DMEM media, which is more relevant for mammalian cells and tissues (**Figure S9b**). To quantify the amount of secreted protein in DMEM, we optimized a split NanoLuc Luciferase assay based on the interaction of two complementary subunits, LgBiT (large bit) and SmBiT (small bit) [33]. The 11-amino-acid SmBiT tag was fused at the C-terminal of elafin, which can bind to the lgBiT fragment added in the culture supernatant to generate a luminescent signal in the presence of furimazine (**Figure 4b**). Based on the luminescence values obtained in this analysis (**Figure 4c**), we correlated the values to a standard curve (**Figure S8a**) and quantified that approximately 30 nM of the elafin_SmBiT fusion protein was detected from culture and PEARL supernatants (**Figure S8b, 4e**). To assess the bioactivity of the secreted elafin in both DMEM culture and PEARLs incubated in DMEM, we performed an in-house optimized neutrophil elastase inhibitory assay (**Figure 4f**, described in materials & methods). The results confirmed that elafin secreted from PEARLs was functional and effectively inhibited neutrophil elastase (NE) (**Figure 4g**).

**Figure 4:**
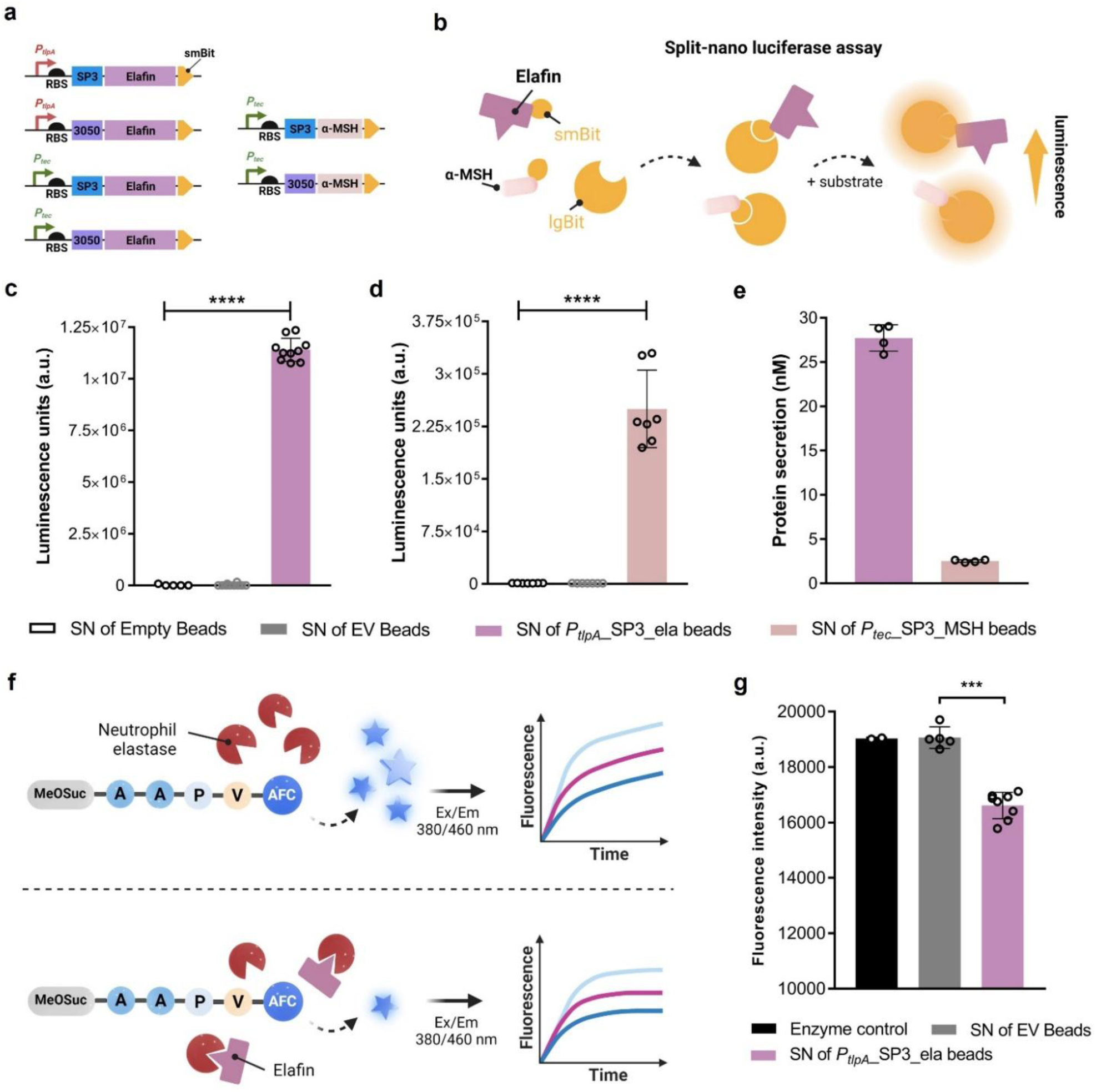
Genetic constructs designed, and assays performed in DMEM media to assess secretion of therapeutically relevant recombinant proteins. **a)** Schematic of genetic constructs of *P_tlpA_* and *P_tec_* promoters driving the expression of elafin and α-MSH, each tagged with either 3050 or SP3 signal peptide and SmBiT fragment. **b)** Graphical illustration of Nano Luciferase assay wherein when LgBiT (large bit) comes in close proximity to elafin_SmBiT or α-MSH_SmBiT, reassembles into a functional luciferase enzyme that produces a luminescent signal upon the addition of a substrate. **c)** A plot representing luminescence values was obtained from a Nano Luciferase assay performed with supernatants of elafin-secreting PEARLs incubated in DMEM media for 24 hours. (n = 10, mean ± SD) **d)** Plot representing luminescence values obtained from Nano Luciferase assay performed with supernatants of α-MSH-secreting PEARLs incubated in DMEM media for 24 hours. (n = 7, mean ± SD) **e)** Quantification of elafin and α-MSH secreted into PEARL supernatants after 24 hours. **f)** Graphical representation of Neutrophil elastase inhibition assay optimized to detect the bioactivity of elafin; Presence of elafin results in decreased fluorescence. **g)** Plot comparing the fluorescence values at 10 minute-time point showing the decreased fluorescence in elafin-PEARL supernatant indicating the NE inhibition by elafin. (n = 5, mean ± SD)

To demonstrate the functionality of the *P_tec_* promoter in secreting a therapeutic protein, we tested two combinations of α-MSH (tagged with Lp_3050 and SP3). α-MSH is an endogenous peptide hormone expressed by keratinocytes in the human body, that stimulates melanogenesis in melanocytes providing photoprotective effects against ultraviolet (UV) radiation among other health benefits. [34] We could not obtain a mutation-free construct for *P_tec_*___3050_α-MSH_SmBiT but could do so for *P_tec_*___SP3_α-MSH_SmBiT. Based on quantification from the split Nano Luciferase assay, 10 nM of α-MSH was secreted by *P_tec_*___SP3_α-MSH_SmBiT construct in DMEM media after 24 hours, and 2.5 nM of α-MSH was detected in the PEARL supernatant.

Depending on the genetic compatibility resulting from the promoter’s strength, the signal peptide’s efficiency, and the protein’s nature, some genetic combinations work efficiently, leading to the secretion of significant amounts of recombinant proteins, while others do not. This highlights the importance of having and testing multiple options of promoters and signal peptides for the production and secretion of different proteins. In all cases where successful protein secretion could be encoded in *L. plantarum*, the PEARL format supports this functionality and release of these differently sized proteins into the surrounding media (α-MSH_SmBiT: 5 kDa, elafin_SmBiT: 11 kDa, NucA: 17 kDa).

### 5) Polycistronic gene expression from PEARLs

Since our strongest promoter, *P_tec_* was capable of driving high levels of NucA secretion and fluorescent protein expression, we tested its ability to do both simultaneously. Being able to encode strong production of two proteins at once would improve the versatility of programming *L. plantarum*. This would be better than encoding separate copies of the promoter for each gene since (i) it reduces the overall size of the genetic module and the resulting plasmid, which is an important parameter affecting transformation efficiency in *L. plantarum*, plasmid retention, and metabolic burden during replication, (ii) eases further genetic manipulation within the plasmid by avoiding the inclusion of repetitive sequences, and (iii) simplifies the regulatory control of both genes at once since only a single promoter needs to be modified to alter their expression levels. For this, we inserted the mScarlet3 gene downstream of the NucA gene, separated by a ribosome binding site (**Figure 5a**). With non-encapsulated bacteria, we observed that the expression of mScalet3 dropped to roughly half when compared to the strain encoding only mScarlet3 production. In contrast, NucA secretion did not significantly decrease compared to the strain only encoding NucA secretion (**Figure 5b**). Both reporters could also be detected well in a DNAse agar plate when grown in static (**Figure 5c**). In the PEARL format, mScarlet3 could be detected inside the bead and continuous secretion over 7 days was also detected using DNAse agar plates (**Figure 5d**).

**Figure 5:**
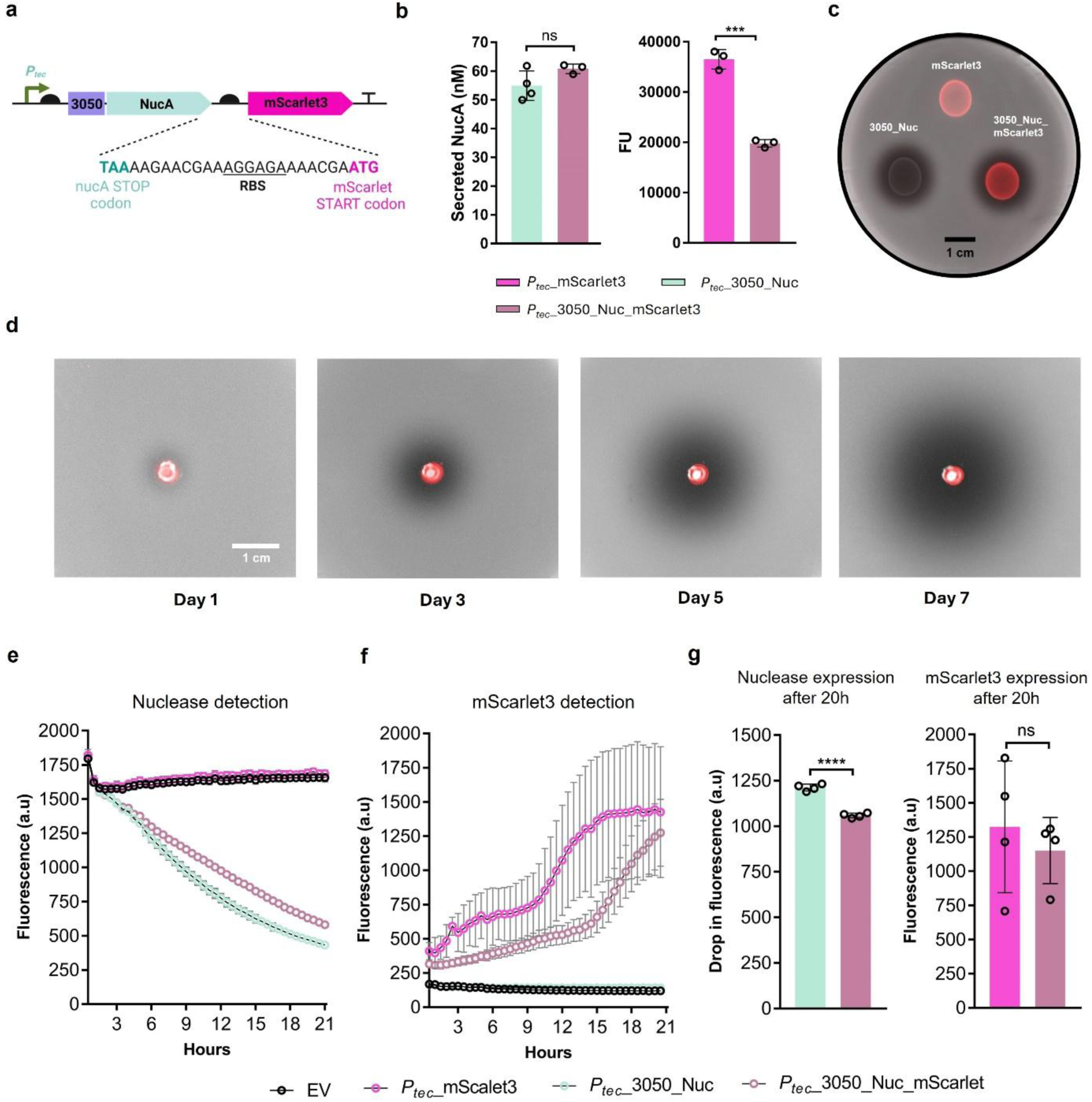
Polycistronic gene expression from PEARLs. **a)** Scheme of the polycistronic gene based on NucA and mScarlet3 genes. The DNA sequence between the stop codon of NucA and the start codon of mScarlet3 is shown. **b)** The left plot shows the levels of secreted NucA (n= 3/4, mean ± SD) for the polycistronic-engineered bacteria and single NucA secreting bacteria (*P_tec_*_3050) when bacteria were grown in DNase media. The right plot shows the decrease in fluorescence expression (mean ± SD) by placing the mScarlet3 gene downstream NucA gene (in comparison to mScarlet3-producing *L. plantarum* WCFS1) when bacteria were grown in MRS media. **c)** DNA agar plate showing bacteria expressing only mScarlet3, secreting only NucA, and expressing and secreting mScarlet3 and NucA, respectively, after 24 hours of growth. **d)** PEARL containing the polycistronic engineer strain. Expression of both genes could be detected from day 1 and kept increasing over the days. **e)** Detection of NucA secretion as drop in fluorescence from PEARLs ( (mean ± SD) incubated in DNase media. **f)** Detection of mScarlet3-driven fluorescence from PEARLs (mean ± SD) incubated in LB media. **g)** Single gene versus polycistronic gene expression levels comparison after 20 hours of growth in PEARLs. Significant differences were only obtained for NucA (left plot).

Furthermore, we assessed the detection of both proteins from the PEARLs by tracking the fluorescence with the microplate reader. We could detect a drop in fluorescence due to NucA secretion (**Figure 5e**) or an increase in mScarlet3-driven fluorescence (**Figure 5f**) only when bacteria encode for NucA or mScarlet3 as a single gene or policystronically. In PEARLs, the differences in protein production after 20 hours of growth were only significant for the NucA (**Figure 5g**). Overall, these experiments confirmed that the *P_tec_* promoter can be used to drive strong expression of two independent genes in parallel, secretion is minimally compromised by doing this, and this functionality is maintained in the PEARLs. Given the lack of incredibly strong constitutive promoters in *L. plantarum* WCFS1, this polycistronic approach supposes an attractive alternative to express two genes of interest highly.

### 6) Cytocompatibility investigation of bacterial alginate gels secreting NucA

*L. plantarum* is known to produce lactic acid and other metabolites that can be cytotoxic to eukaryotic cells. [39] Several reports suggest that such cytotoxicity can be selective for cancer cells but we found it important to test if encapsulation of these bacteria in the PEARLs would alter this cytotoxicity. To evaluate this, we co-cultured the PEARLs along with human fibroblasts in a media optimized to support the growth of *L. plantarum* and human cells. Thus, during the 24-hour period of this experiment, the bacteria proliferate within the PEARLs and secrete NucA. These co-cultures were used to assess the impact of PEARLs on the viability and metabolic activity of human fibroblasts (**Figure 6a**). We compared conditions containing the *P_tec_*_3050 (a high NucA producer, >25 nM, **Figure 3f**) and *P_tlpA_*_3050 (a low NucA producer, <5 nM, **Figure 3f**) strains. We used empty alginate macro beads, wild-type lactobacilli entrapped in the alginate macro beads, NucA-laden alginate macro beads (50 nM), and media containing 0.5, 5, and 50 nM of NucA as controls. First, we quantified the percentage of cell death by measuring the number of cells with damaged cell membranes using the lactate dehydrogenase assay (**Figure 6b**). A lysis condition where all cells were damaged with Triton X-100 was used as a positive control. All conditions investigated presented similar values of cell death around 10%, which were comparable to the negative control, untreated cells. Cell viability was also quantified as the number of metabolically active cells using the alamar Blue assay (**Figure 6c**). Results show similar values of viability to the negative control. The metabolic activity of fibroblasts decreased slightly for *P_tlpA_*_3050, but this was not statistically significant compared to the negative control. Some conditions presented slightly higher metabolic activity than the negative control, which was observed for conditions containing bacteria and without bacteria (i.e., Nuc control 5nM). This points to a different proliferation rate unrelated to the presence of bacteria in the co-cultures. All measurements were within a similar range.

**Figure 6:**
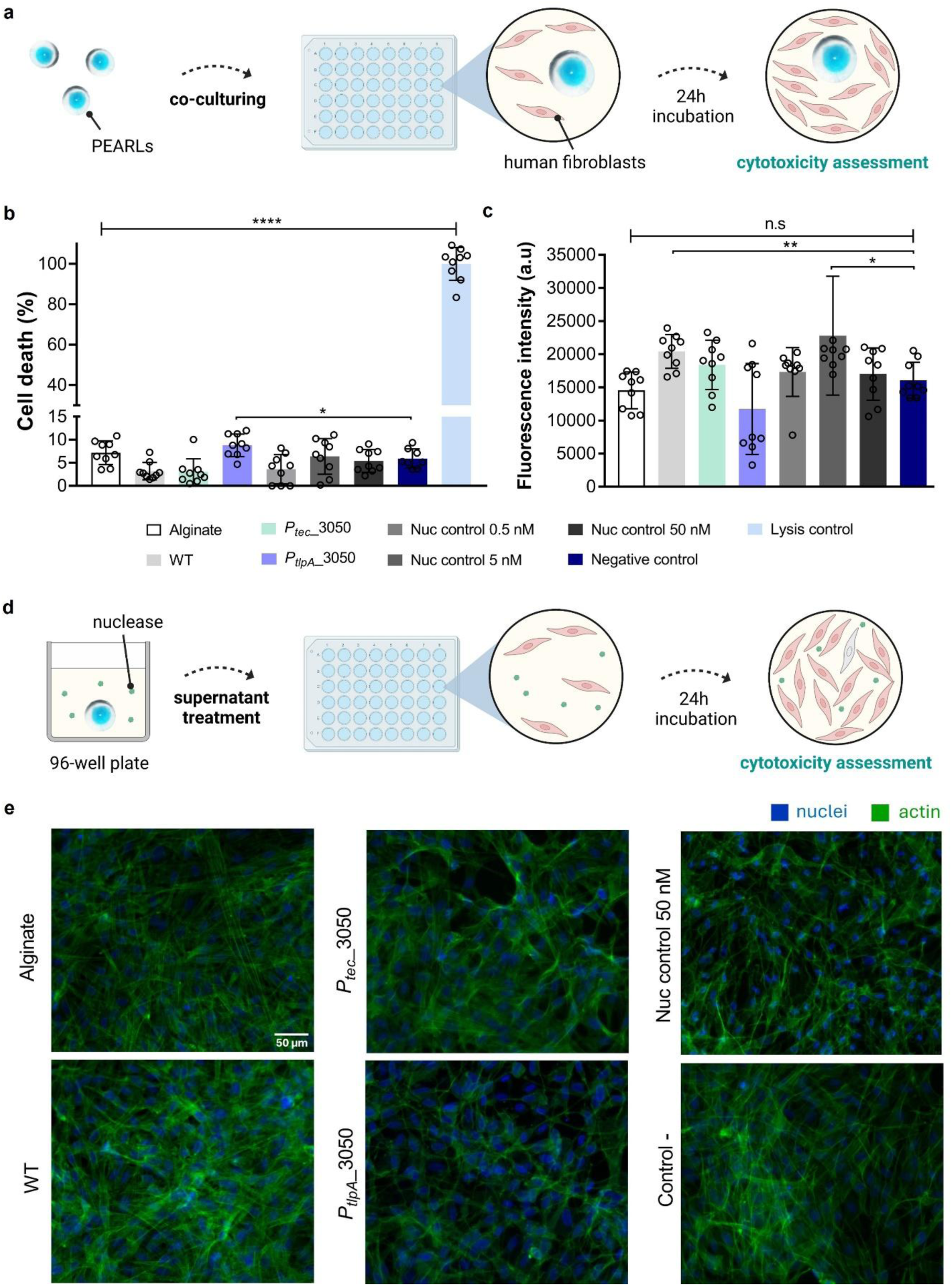
Cytocompatibility evaluation of NucA-secreting PEARLs on human fibroblasts. **a**) Schematic showing PEARLs were co-cultured along with human fibroblasts in growth media (for fibroblasts) for 24 h to evaluate the compatibility potential on human fibroblasts. The collected supernatants were used viability assessment. **b**) Percentage of cell death quantified after co-culture treatment via LDH assay (mean ± SD). **c**) Fluorescence intensity measured during alamarBlue assay after co-culture treatment (mean ± SD, arbitrary units (a.u)). **d)** Schematic showing PEARLs incubation in growth media (for fibroblasts) overnight for evaluation by immunostaining. The collected supernatants were used to treat fibroblasts in culture for 24 h, after which immunostaining was performed. **e**) Representative fluorescence images of human fibroblasts after treatment with PEARLs supernatants (scale bar: 50 μm, blue: nuclei, green: actin).

LDH assay and alamar blue assay were performed with PEARL supernatants as well and the results show similar values as negative control (**Figure S10a-b**). We also stained the actin cytoskeleton of the fibroblasts with supernatants of PEARLS (**Figure 6d**) to assess cell morphology (**Figure 6e**). From the staining, we observed similar morphologies across conditions, with spread cells and mature actin fibers developed. Nuclei staining was quantified to obtain values of cell densities for the conditions evaluated. Cell densities were similar across conditions (∼1600 cells/mm^2^) indicating comparable cell confluency after treatment to the negative control **(Figure S11**).

### 7) Zebrafish Embryo toxicity test

Zebrafish (ZF) embryos <5 days post fertilization (dpf) were used as a highly sensitive *in vivo* acute toxicity invertebrate test model. Following OECD guideline 236 [41], such tests are usually performed in 24-well plates with one ZF embryo per well in a volume of 2 mL media. In order to achieve a more robust exposure scenario, the test protocols were adapted in the following ways to be better suited for PEARLs: Usually, the zebrafish embryos are exposed to test compounds while they remain inside the egg from which they hatch later during the test. However, the chorion can prevent substances from accessing the embryo. [42] To ensure that the embryos got into direct contact with any bacterial products released in the media, they were released from the chorion before the start of the test. The de-chorionation took place at 1 dpf since this is the earliest time at which this treatment can be carried out reliably without harming the embryos, and which is also a common practice in drug screening. [43]

The ZF media was adapted to support *L. plantarum* growth and metabolism by adding saccharose. Different sugars (Glucose, Galactose, Maltose, Arabinose, Fructose, Sorbitol, Xylose, Mannose and Saccharose) were added to the *L. plantarum* culture at 1% v/v concentration and after 24 hours, OD_600_ and pH of the cultures were measured. Saccharose supplementation was the only condition in which significant bacterial growth (up to OD_600_ of 2) was seen along with the least drop in pH to 6. In all the remaining conditions with other sugars, either bacterial growth was compromised, or the pH decreased below 6, even down to 3.

The potential toxicity of non-encapsulated *L. plantarum* on ZF embryos was investigated using a cell-free supernatant containing the complete bacterial secretome. To decrease the production volumes needed, the test was performed in a 96-well plate, thereby decreasing the test volume from 2 to 0.3 mL per replicate. We observed that the pure supernatant instantly killed all ZF embryos within the first 24 hours (**Figure 7b**). On dilution to 50 % of the culture supernatant, 45 % of the ZF embryos died within the first 24h. After 48 hours all embryos were dead. On dilution to 25 % of the culture supernatant, 75 % survived after 24 h, 55 % after 48 h, and 50 % after 72 and 96 h. The results indicate that metabolic byproducts of *L. plantarum* are toxic to ZF embryos, similar to their reported effect on cancer cells. [39] Notably, the supernatant decreased the pH from 8 to values around 6, possibly due to lactic acid production by the bacteria. This low pH was measured in the 50% diluted samples but in 25% dilution, the pH was closer to neutral (**Table S2**). However, ZF embryos have a relatively high pH tolerance and should tolerate the observed pH changes. [44] Therefore, the observed mortality induced by the supernatant is considered to be induced rather by substances produced by *L. plantarum* than by decreased pH.

**Figure 7:**
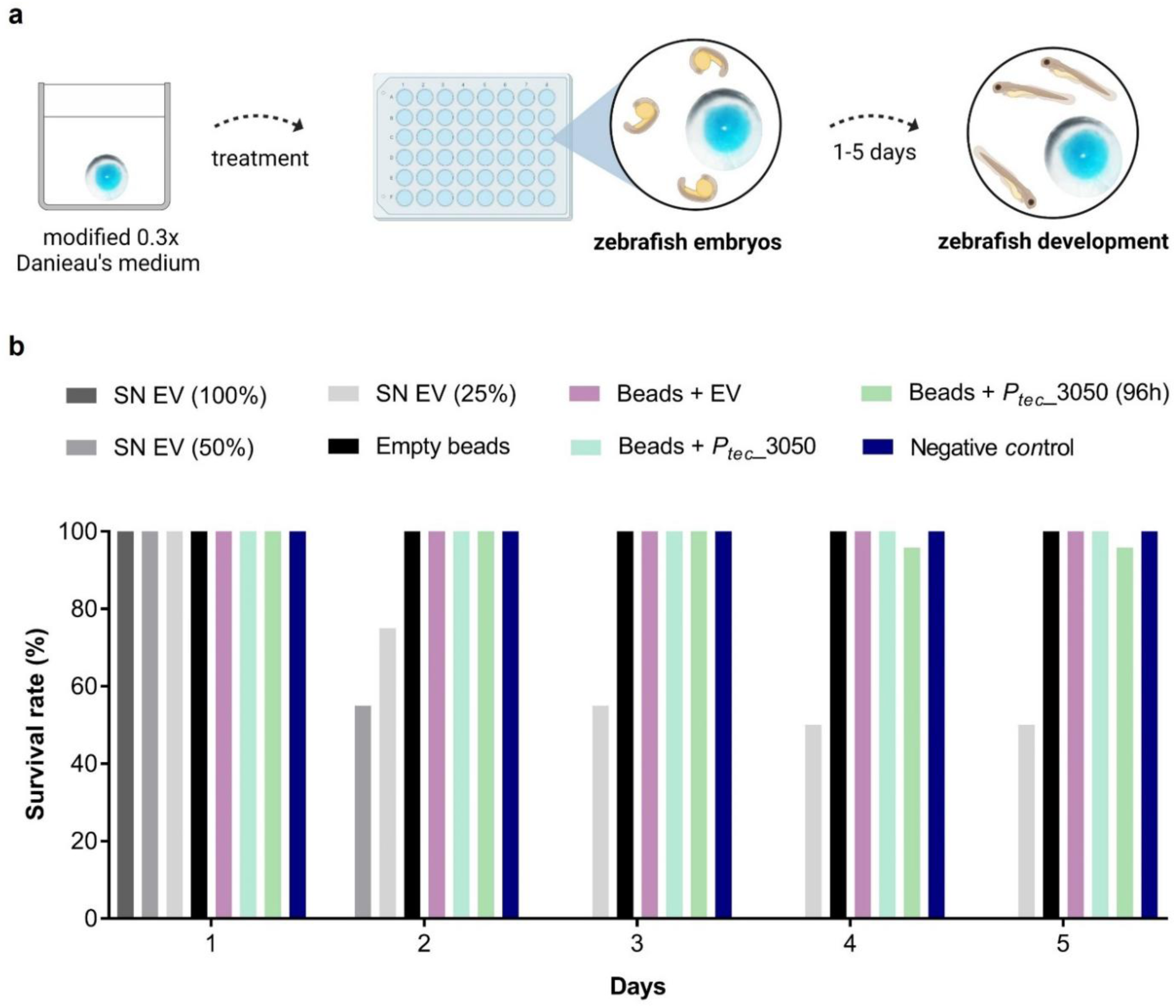
Cytotoxicity evaluation of NucA-secreting alginate beads upon Zebrafish embryos. **a)** Graphical representation of alginate beads incubated in 0.3x Danieu’s media followed by incubation of zebrafish embryos with beads in a 48-well flat bottom plate up to 5 days. **b)** Survival rates of zebrafish embryos exposed to three different concentrations of the cell-free supernatant (SN) containing the complete bacterial secretome (SN EV (100%), SN EV (50%), SN EV (25%) as a positive control (n=20/treatment group), four different kinds of alginate beads: empty beads, beads + EV (containing wild-type *L. plantarum*), beads + *P_tec_*_3050 ( secreting the highest level of NucA), beads + *P_tec_*_3050 (96h) (pre-incubated for 96h before adding ZF embryos) and the negative control (n=24/treatment group).

The effect of the PEARLs on ZF embryos were tested in 48-well plates to reduce the volume and increase the concentration of potentially released bacterial metabolic byproducts. The size of a well was small enough to establish direct contact between embryos and beads, but still big enough to prevent mechanical interferences such as squeezing of the embryos. 8 PEARLs were tested for each condition with 3 embryos in the same well as one PEARL. The scheme of the experiment is shown in **Figure 7a**.

Survival of the ZF embryos in the presence of the PEARLs was tested over 96 hours and compared to a negative control group that did not contain any PEARLs or alginate. In the first test group, empty alginate beads without bacteria were tested. In the second group, core-shell alginate beads contained wild-type *L. plantarum*. The next group included PEARLs containing the *P_tec_*___3050 strain, which secretes the highest level of NucA. In all these groups, all embryos survived the 96-hour incubation period (**Figure 7b**). In an additional treatment group, the PEARLs containing the *P_tec__*3050 strain were incubated for 96 hours without ZF before the start of the test to increase the concentration of secretion products in the surrounding media. In this treatment, one embryo died between test day 3 and 4 resulting in a survival rate of 95.8 % (**Figure 7b**). Such effects with a value lower than 10 % are usually considered natural variation and are not related to toxicity [40]. Even though a toxic effect cannot be completely ruled out, the death of a single embryo is attributed to natural variance rather than a toxic effect, supported by the absence of any other adverse effects. Thus, no significant reduction of survival due to the presence of PEARLs was observed during the 96h test duration. This was quite a surprising result considering the toxicity of the non-encapsulated bacteria. It is to be noted that the PEARLs did not considerably reduce the pH of the media as it remained around 7.5 in all cases (**Table S2**). This indicated that encapsulation of the bacteria lowered its lactic acid production rate. While it is possible that alginate encapsulation slowed down metabolite diffusion into the media, the results from **Figure 3d** indicate that diffusion of a secreted protein occurs within a matter of hours, so the 4-8 day periods of the last two test groups should have been sufficient for metabolite accumulation in the media. Furthermore, we were able to clearly detect NucA secretion into the ZF media after 24 hours (**Figure S12a**).

In parallel, the PEARLs incubated in ZF media for 14 days were dissolved using EDTA to recover the viable bacteria and plated on an MRS-agar plate. The number of colony-forming units (CFU) was found to be around 10^5^ CFU/mL (**Figure S12c**). Though this is lower than the initial number of cells that were encapsulated in the PEARLs (∼10^6^), it is to be noted that EDTA treatment could have killed some of the bacteria. [27] We confirmed this by dissolving PEARLs 1 hour after their fabrication and determining the bacterial counts using the same method, which was in the same order of magnitude (<4×10^5^ CFU/mL). However, this was still higher than the 14-day-old PEARLs, so it is possible that bacterial viability or robustness dropped over this time period. Additionally, six random colonies from the MRS agar plate were spotted on the DNase-agar plate, and secretion was observed from all colonies after 24 hours (**Figure S12d**). These results indicated that PEARLs could maintain the viability and activity of a bacterial population over 14 days and that encapsulation of *L. plantarum* in alginate possibly reduces or prevents the production of metabolites that can be cytotoxic to the zebrafish. Extensive analysis of the secretome from non-encapsulated bacteria and the PEARLs using metabolomic and proteomic methods could shed light on this mechanism.

## CONCLUSIONS

To our knowledge, this is the first study that comprises an ELM based on probiotic lactobacilli. We have developed a rapid, cost-effective, and safe encapsulation method that keeps probiotic bacteria viable and active while containing them. On top of that, the manual approach of fabricating PEARLs is cost-effective and easy to implement for small-batch production at the lab scale. Characterization of these PEARLs has revealed interesting insights concerning protein secretion and cytotoxicity. For example, the exact amount of secreted protein from both non-encapsulated and encapsulated bacteria was measured in the range of 5 – 50 nM and, a majority of this was secreted within the first 6h. Encapsulation as PEARLs offered multiple benefits like supporting long-term protein release (at least 14 days), stabilizing the release profiles, and improving release rates. Additionally, we discovered and characterized new genetic parts that will enhance the programmability of *L. plantarum* for protein secretion. Two constitutive promoters, *P_tec_* and *P_tlpA_* secreted significant amounts of three different proteins (NucA, elafin, and α-MSH) from PEARLs. Finally, the cytotoxicity results using human cell lines and ZF suggest considerable suppression of toxic metabolite release when *Lactobacilli* are encapsulated in alginate. Altogether, these results indicate that PEARLs are a promising platform for further development towards therapeutic applications.

## SUPPORTING INFORMATION

A file containing all the supporting information is available from the Wiley Online Library.

## ACKNOWLEDGEMENTS

We thank Prof. Wilfried Weber for generously sharing the genetic parts encoding LgBiT and SmBiT, and Anwesha Chatterjee for assisting with the Nano Luciferase assay. We also thank Dr. Cao Nguyen Duong for his help and support with the microscopy experiments. We thank Therese Steudter for providing the Alcian blue dye. The pLp_3050sNuc plasmid was a kind gift from Prof. Geir Mathiesen (Addgene plasmid # 122030). We thank Dr. Samuel Pearson for suggesting α-MSH peptide as a therapeutic for ELMs. Biorender was used for generating the schemes in this manuscript.

## DATA AVAILABILITY

The information provided in the main text and supporting information are sufficient to reproduce the experiments described in this study. The raw and processed datasets along with relevant metadata are available from the corresponding author upon reasonable request.

