## Supplementary information for "PEARL: Protein Eluting Alginate with Recombinant Lactobacilli"

[†] Joint Authors

*

**Table 1**. List of the primers used in this study

| Primer name | Sequence |
|---|---|
| **Vector mScarlet3/Nuc/Ela/α-MSH fw** | 5'- gaagataaatcccataaggg -3' |
| **Vector mScarlet3/Nuc/Ela/α-MSH rev** | 5'- aagaactctattgaagccc -3' |
| **Vector Operon fw** | 5'-aataactagcataacccc-3' |
| **Vector Operon rev** | 5'-ttattgacctgaatcagcg-3' |
| **Insert mScarlet3 Operon fw** | 5'-acgctgattcaggtcaataaaagaacgaaaggagaaaac -3' |
| **Insert mScarlet3 Operon rev** | 5'-ggggttatgctagttatt-3' |
| **Direct Protocol mScarlet3 fw** | 5'-gatcgttcgtagagatcg-3' |
| **Direct Protocol mScarlet3 rev** | 5'-cttgctcaacttgtggtt-3' |
| **Direct Protocol Nuc/ Ela/α-MSH/Operon fw** | 5'-ccgttagcgtagtagtagc-3' |
| **Direct Protocol Nuc/ Ela/α-MSH/Operon rev** | 5'-cttcaaagggtcaacagc-3' |

- **SP1_Nuc**

**MKLLLVLLQQLLVHFEASSKALLCK**HHHHHHPLSTKKLHKEPATLIKAIDGDTVKLMYKGQPMT
FRLLLVDTPETKHPKKGVEKYGPEASAFTKKMVENAKKIEVEFDKGQRTDKYGRGLAYIYADGK
MVNEALVRQGLAKVAYVYKPNNTHEQHLRKSEAQAKKEKLNIWSEDNADSGQ

- **SP2_Nuc**

**MQKKTVGLLLASAGPFLWGSSGTVAQHLF**HHHHHHPLSTKKLHKEPATLIKAIDGDTVKLMYK
GQPMTFRLLLVDTPETKHPKKGVEKYGPEASAFTKKMVENAKKIEVEFDKGQRTDKYGRGLAYI
YADGKMVNEALVRQGLAKVAYVYKPNNTHEQHLRKSEAQAKKEKLNIWSEDNADSGQ

- **SP3_Nuc**

**MKKIVNWLLGSVLMIAAVTMLSSVSANASTYY**HHHHHHPLSTKKLHKEPATLIKAIDGDTVKLM
YKGQPMTFRLLLVDTPETKHPKKGVEKYGPEASAFTKKMVENAKKIEVEFDKGQRTDKYGRGL
AYIYADGKMVNEALVRQGLAKVAYVYKPNNTHEQHLRKSEAQAKKEKLNIWSEDNADSGQ

- **SP4_Nuc**

**MKKFMNSNWSMRLLALLCALVLFAYVNSLKTT**HHHHHHPLSTKKLHKEPATLIKAIDGDTVKLM
YKGQPMTFRLLLVDTPETKHPKKGVEKYGPEASAFTKKMVENAKKIEVEFDKGQRTDKYGRGL
AYIYADGKMVNEALVRQGLAKVAYVYKPNNTHEQHLRKSEAQAKKEKLNIWSEDNADSGQ

- **SP5_Nuc**

**MKKWSTYNKNTSHKIVELSSIVVMSSVLGIILAKPVQGKADQTS**HHHHHHPLSTKKLHKEPATL
IKAIDGDTVKLMYKGQPMTFRLLLVDTPETKHPKKGVEKYGPEASAFTKKMVENAKKIEVEFDK
GQRTDKYGRGLAYIYADGKMVNEALVRQGLAKVAYVYKPNNTHEQHLRKSEAQAKKEKLNIWS
EDNADSGQ

- **Lp_3050_Nuc**

**MKKFNFKTMLLLVLASCVFGVVVNVTTSLGPQTAITAQASKKL**HHHHHHPLSTKKLHKEPATLI
KAIDGDTVKLMYKGQPMTFRLLLVDTPETKHPKKGVEKYGPEASAFTKKMVENAKKIEVEFDKG
QRTDKYGRGLAYIYADGKMVNEALVRQGLAKVAYVYKPNNTHEQHLRKSEAQAKKEKLNIWSE
DNADSGQ

- **Lp_3050_ela_smBit**

**MKKFNFKTMLLLVLASCVFGVVVNVTTSLGPQTAITAQASKKL**EAAVTGVPVKGQDTVKGRVP
FNGQDPVKGQVSVKGQDKVKAQEPVKGPVSTKPGSCPIILIRCAMLNPPNRCLKDTDCPGIKK
CCEGSCGMACFVPQGGGGSGGGGSVSGWRLFKKIS

- **SP3_ela_smBit**

**MKKIVNWLLGSVLMIAAVTMLSSVSANASTYY**EAAVTGVPVKGQDTVKGRVPFNGQDPVKGQ
VSVKGQDKVKAQEPVKGPVSTKPGSCPIILIRCAMLNPPNRCLKDTDCPGIKKCCEGSCGMAC
FVPQGGGGSGGGGSVSGWRLFKKIS

- **Lp_3050_α-MSH_smBit**

**MKKFNFKTMLLLVLASCVFGVVVNVTTSLGPQTAITAQASKKL**HHHHHHGGSGGSVSGWRLF
KKISGGSGGSSYSMEHFRWGKPV

- **SP3_α-MSH_smBit**

**MKKIVNWLLGSVLMIAAVTMLSSVSANASTYY**VSGWRLFKKISGGGGSGGGGSHHHHHHGG
GGSGGGGSSYSMEHFRWGKPV

**Figure S1**. Protein sequences from all the secretion modules used in this study. Signal peptides are shown in bold and red. The histidine tag is shown in purple. NucA sequence is shown in green. Elafin sequence is shown in pink. α-MSH sequence is shown in yellow. smBit sequence is shown in orange. Linkers are shown in blue.

- ***$P_{tlpA}$* promoter**

TTTAATTTGTTTGTTAGTTAGTTTATTTGTTGGTTTGTTTGTGTTATAATATCCTCTAGA
AATAATTTTGTTTAACTTTAAGA**AGGAGG**TATACCATG

- ***$P_{tec}$* promoter**

ACAAAAAGTTGAAAAAAGTTGACTTTTTGTGTTGACTCCCGGTCTAGAACGGGGTAT
TATTAAAGCATCAAGAACGAA**AGGAGA**AAACGAA

**Figure S2**. DNA sequences of the promoters used in this study. Ribosome-binding sites are shown in bold.

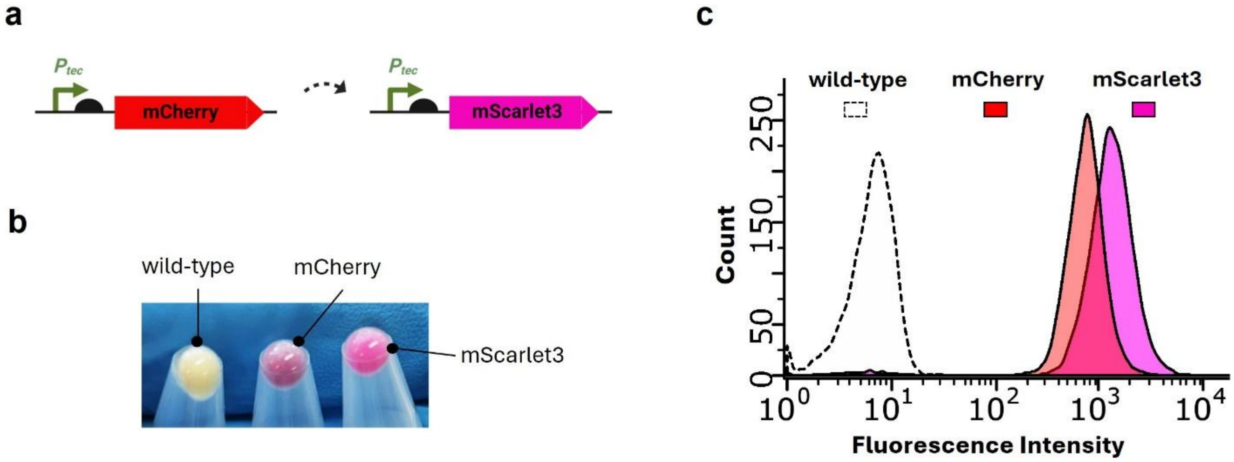


**Figure S3**. **a)** Scheme of the reporter change from mCherry to mScarlet3. **b)** Bacterial pellets from wild-type *L. plantarum* WCFS1, *L. plantarum* WCFS1 producing mCherry and *L. plantarum* WCFS1 producing mScarlet3. **c**) Flow cytometry plot showing the fluorescence intensity of wild-type *L. plantarum* WCFS1, *L. plantarum* WCFS1 producing mCherry and *L. plantarum* WCFS1 producing mScarlet3.

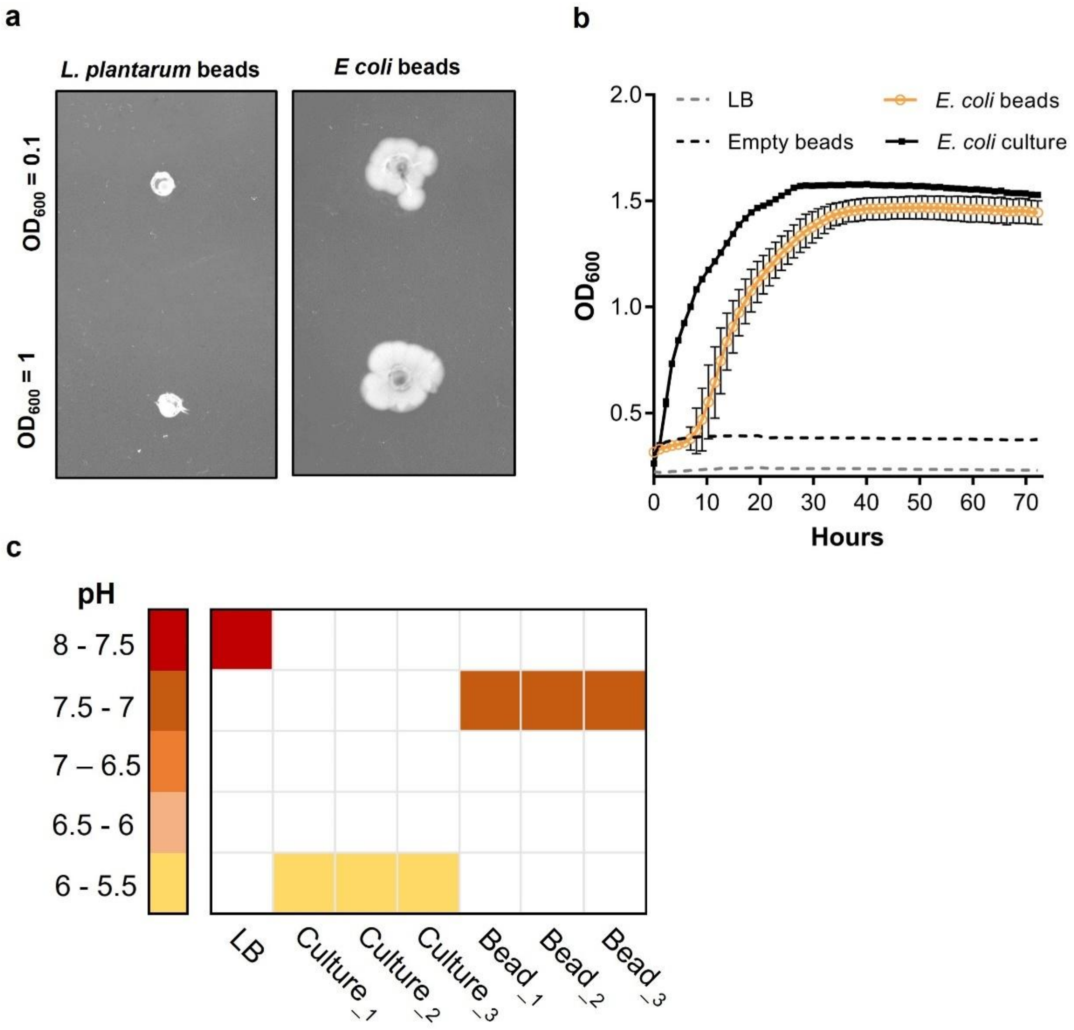


**Figure S4**. **a)** Alginate beads containing either wild-type *L. plantarum* or *E. coli* Nissle at initial OD$_{600}$ of 0.1 and 1 placed on an LB-kanamycin agar plate (for *E. coli* Nissle) and MRS-erythromycin agar plate (for *L. plantarum)* after 24 hours incubation. **b**) 72-hour growth kinetic measurements depicting OD$_{600}$ values of encapsulated *Escherichia coli Nissle* in PEARLs and non-encapsulated culture control (n=4, mean ± SD). **c)** pH values, measured with pH paper strips, from the supernatants (in LB media) of *L. plantarum* cultures and *L. plantarum* beads after 24 hours of growth at 37°C. Experiments were performed as triplicates (n = 3)

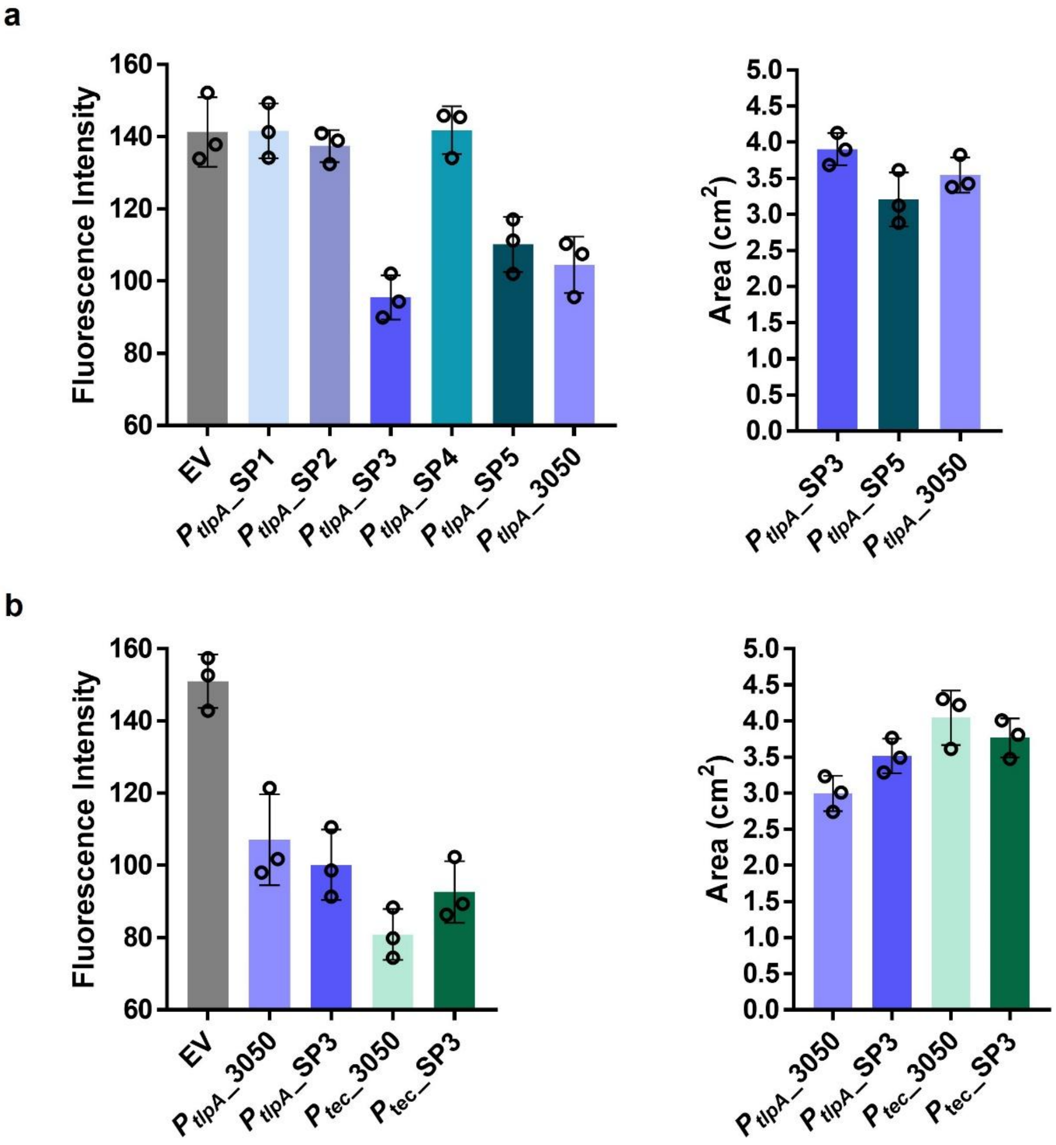


**Figure S5**. **a)** The left plot shows the decrease in fluorescence intensity caused by the discoloration of the DNA agar plate due to the halo formation of each NucA clone (n=3, mean ± SD). The right plot shows the area in $cm^2$ of the halos formed in the DNA agar plate by the three clones that secreted NucA (n=3, mean ± SD). **b)** The plot on the left shows the decrease in fluorescence intensity by each promoter/signal peptide combination (n=3, mean ± SD). The right plot shows the area in cm2 by each each promoter/signal peptide combination (n = 3, mean ± SD).

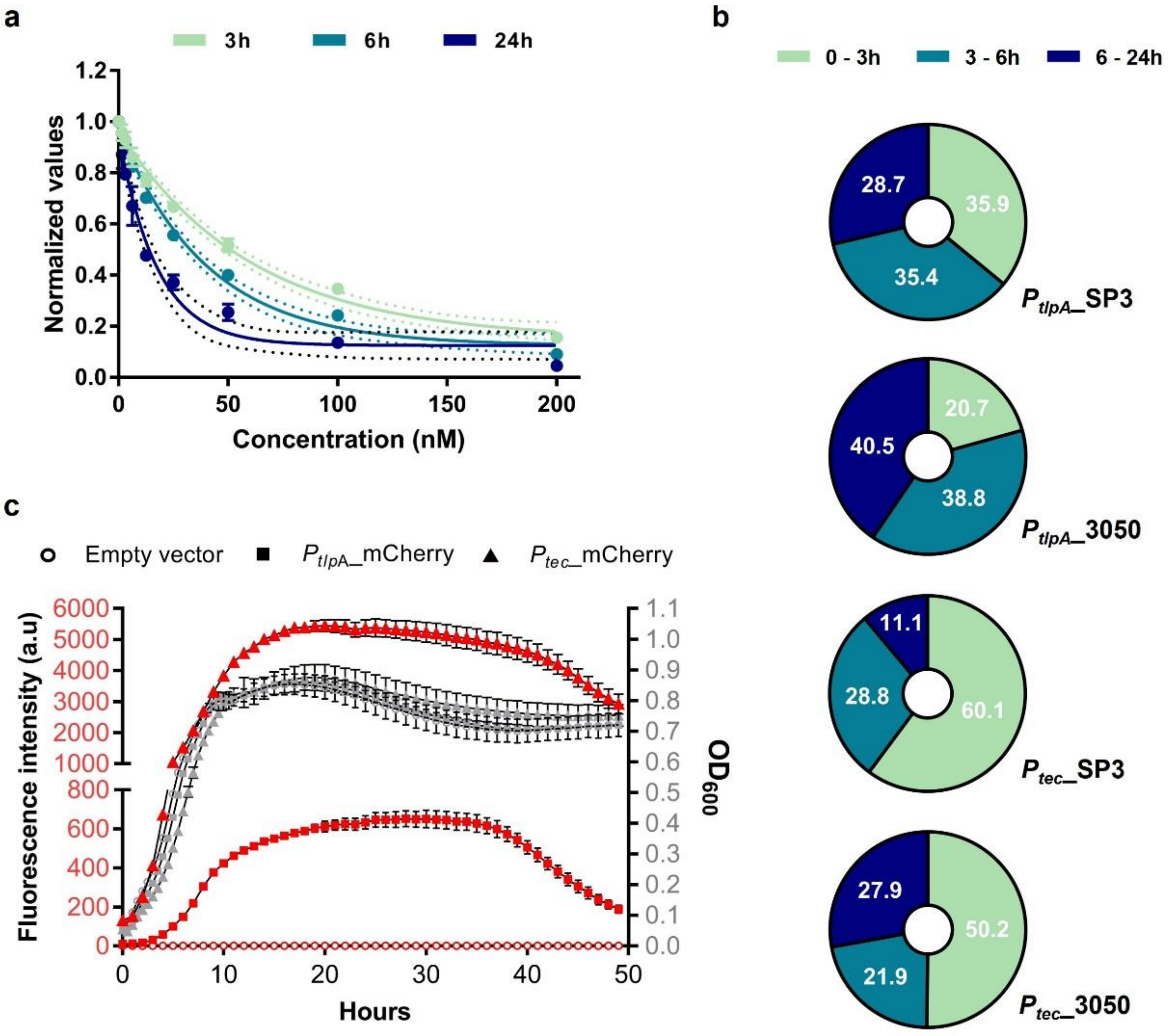


**Figure S6**. **a)** Standard curve for pure NucA after 3, 6 and 24 hours of incubation at 37 °C in shaking conditions (n=3, mean ± SD). **b)** Pie charts showing the percentage of the total NucA secreted after 3, 6 and 24 hours by each promoter/signal peptide combinations. **c)** Plot showing both the differences in the levels of fluorescence (i.e. promoter activity) and growth (as $OD_{600}$) in MRS media for empty vector *L. plantarum*, $P_{tlpA}$_mCherry *L. plantarum* and $P_{tec}$_mCherry *L. plantarum* (n=3, mean ± SD).

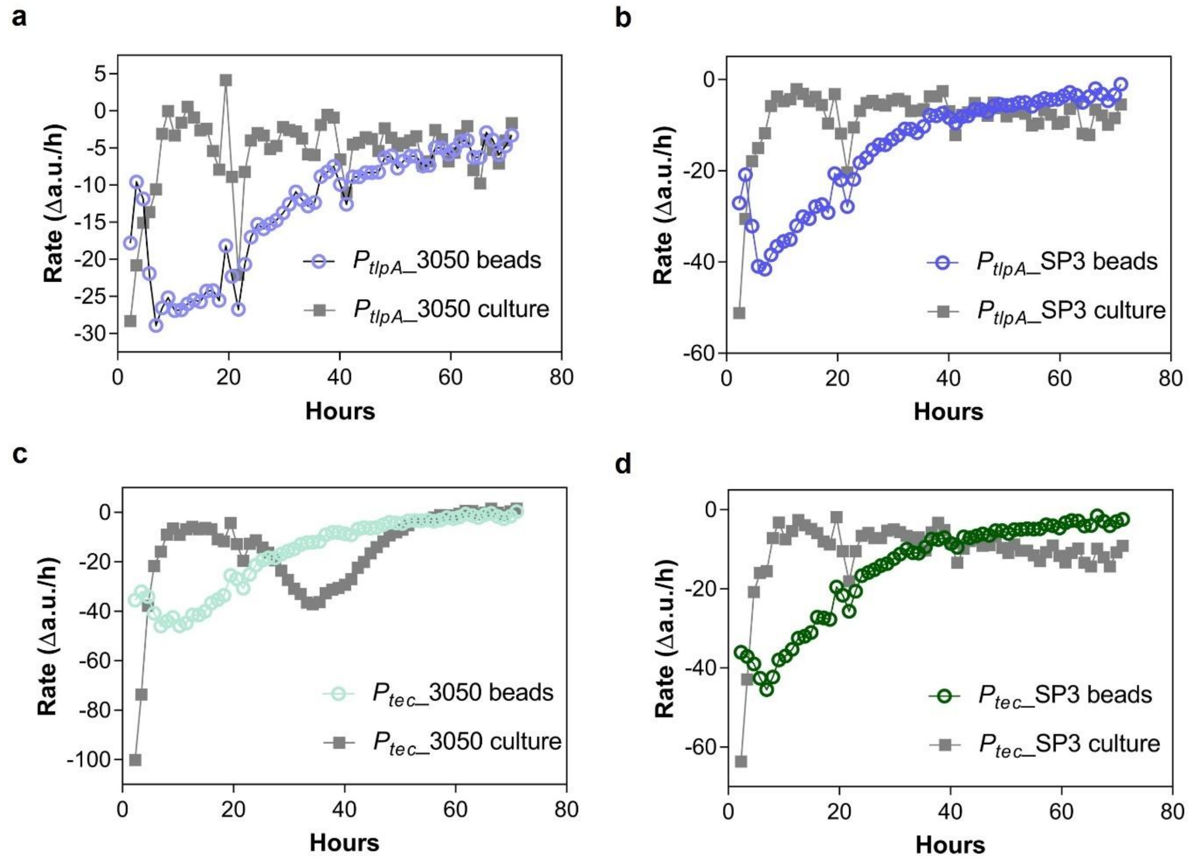


**Figure S7**. **a)** NucA secretion rate by $P_{tlpA}$_3050 in culture and within PEARLs. **b)** NucA secretion rate by $P_{tlpA}$_SP3 in culture and within PEARLs. **c)** NucA secretion rate by $P_{tec}$_3050 in culture and within PEARLs. **d)** NucA secretion rate by $P_{tec}$_SP3 in culture and within PEARLs. All the experiments were performed as experimental quadruplicates (in culture) (n = 4) or quintuplicates (in beads) (n = 5).

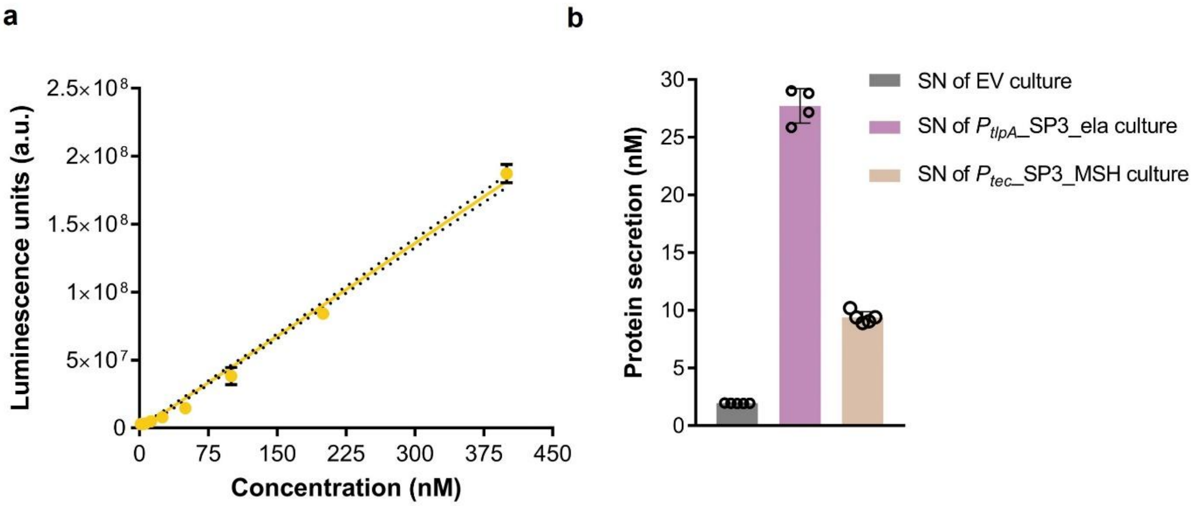


**Figure S8**. **a**) Standard curve of smBiT peptide and LgBiT (1 µM) with furimazine substrate. **b**) Quantification of elafin and α-MSH secreted by non-encapsulated bacteria incubated in DMEM media after 24 hours.

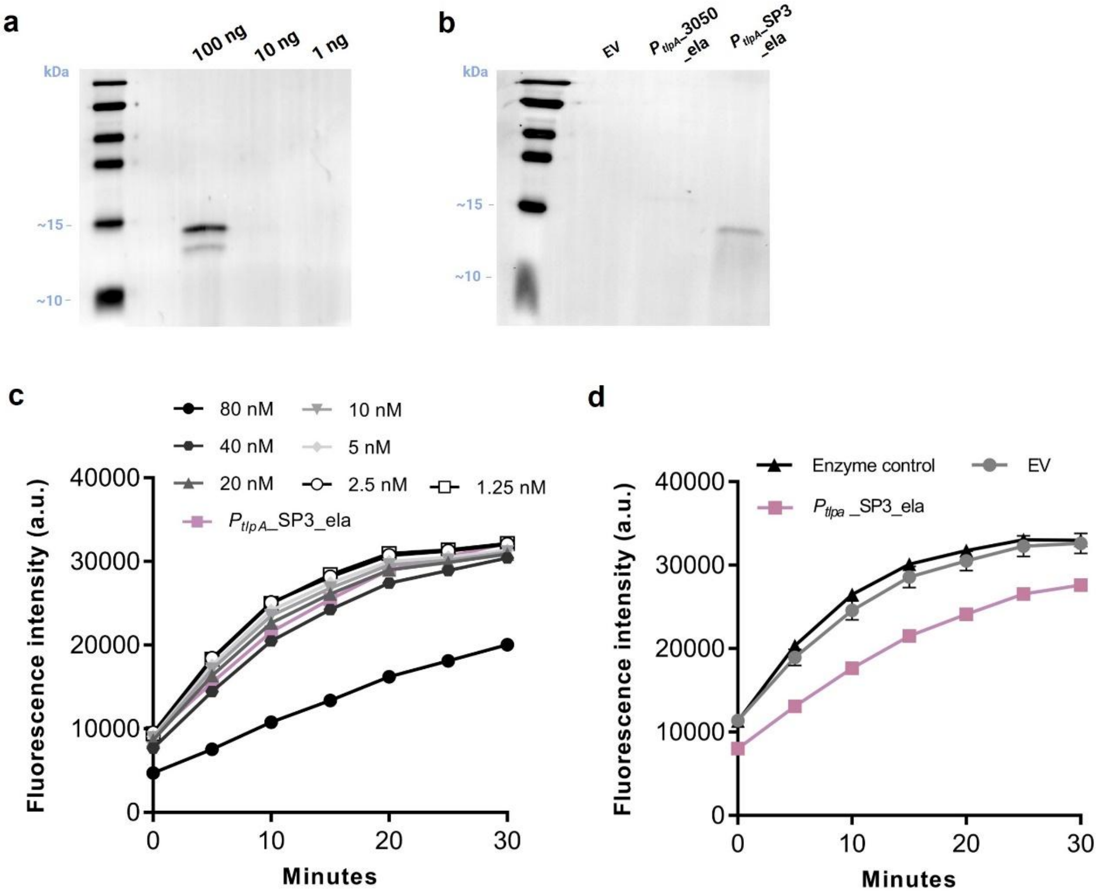

**Figure S9**. Elafin quantification and bioactivity assessment: a) Western blot of difference concentrations of commercial elafin (purchased from R&D Biotechne GmbH) to check the sensitivity, using anti-elafin H2 antibody (Alexa fluor-647) (Santa Cruz Biotechnology) and Spectra Multicolor Broad range protein ladder (Cat 26634, Thermo Scientific). b) Western blot of culture supernatants of wcfs1_EV, wcfs1_$P_{tlpa}$_3050_ela, and wcfs1_$P_{tlpa}$_sp3_ela after incubation in DMEM at 37 °C incubator for 24 hours. c) Time-based kinetics of Neutrophil elastase inhibition (NE) by different concentrations of commercial elafin in DMEM media in the presence of fluorescent substrate. (n=3) D) Time-based kinetics of NE inhibition by culture supernatants of wcfs1-EV and wcfs1_$P_{tlpa}$_sp3_ela, after incubation in DMEM media at 37 °C incubator for 24 hours; Significant decrease in the fluorescence of wcfs1_$P_{tlpa}$_sp3_ela culture supernatant in comparison to wcfs1_EV and enzyme control (negative control) confirms the bioactivity of elafin.

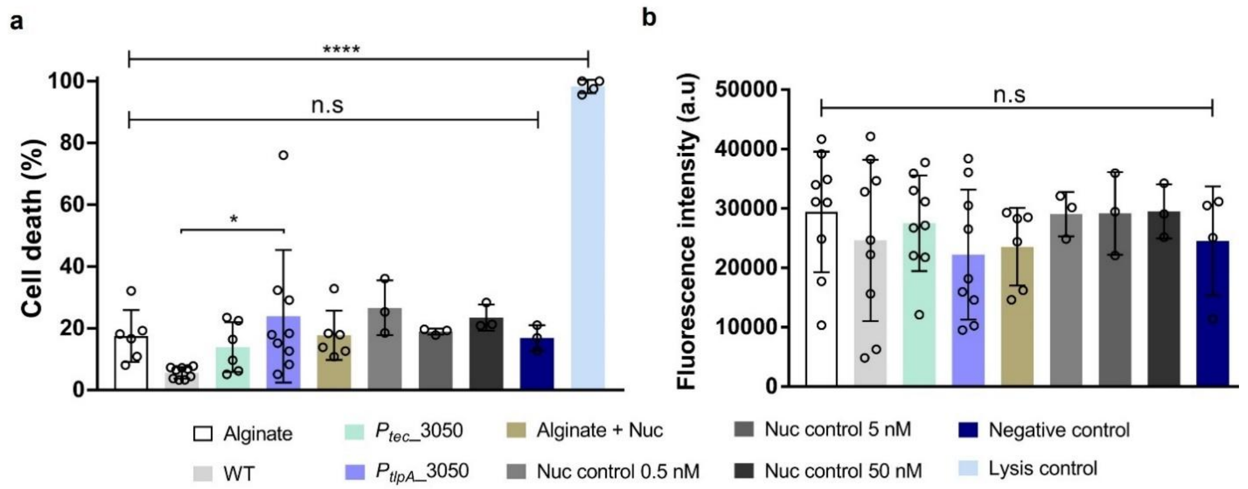


**Figure S10**. **a**) Percentage of cell death quantified after supernatant treatment via LDH assay (mean ± SD). **b**) Fluorescence intensity measured during alamarBlue assay after supernatant treatment (mean ± SD, arbitrary units (a.u)).

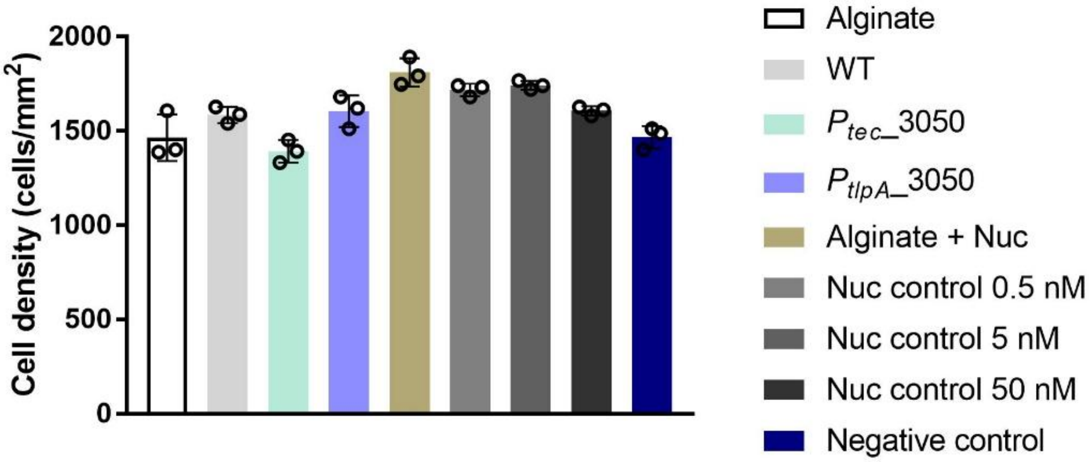


**Figure S11**. Plot showing the cell densities (as cells/mm$^2$) (n=3, mean ± SD) of human fibroblasts obtained 24 hours after the supernatant treatments for all the different conditions.

**Table S2**. pH values from the different conditions assessed during the zebrafish embryo toxicity tests.

| Supernatant Test | n | mean pH | ± SD |
|---|---|---|---|
| **Negative control** | 20 | 7.5 | 0.107 |
| **SN EV (100 %)** | 20 | 6.1 | 0.110 |
| **SN EV (50 %)** | 20 | 6.0 | 0.188 |
| **SN EV (25 %)** | 20 | 6.9 | 0.604 |

| Alginate Beads Test | n | mean pH | ± SD |
|---|---|---|---|
| **Negative control** | 8 | 7.6 | 0.005 |
| **Empty Beads** | 8 | 7.5 | 0.046 |
| **Beads + EV** | 8 | 7.6 | 0.045 |
| **Beads + $P_{tec}$_3050** | 8 | 7.5 | 0.021 |
| **Beads + $P_{tec}$_3050 (96h)** | 8 | 7.5 | 0.035 |

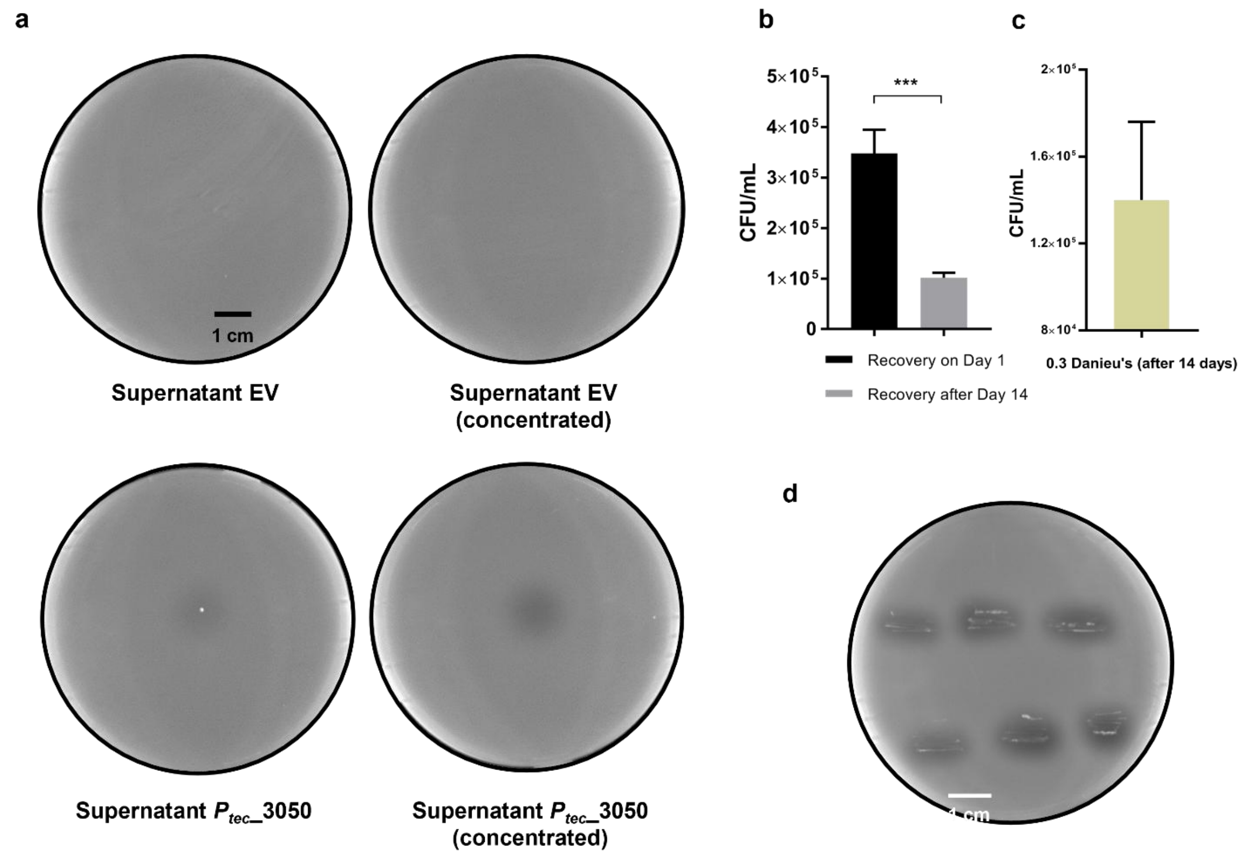


**Figure S12**. **a)** DNA agar plates showing NucA secretion in ZF medium by EV and $P_{tec}$_3050-Nuc secreting bacteria. 15 µL of the SNs of bacteria $OD_{600}$ of 1 (growing in ZF medium for 24 hours) were spotted in DNA agar plates and incubated at 37 °C for 24 hours. The SN were also concentrated, and 15 µL were spotted in DNA agar plates and incubated in the same conditions. **b)** Plot showing recovery of bacteria (wcfs1 $P_{tec}$_3050 nuc) from two sets of PEARLs incubated in LB medium. One set (n=3, mean as column height) of PEARLs were lysed and recovered using 25 mM EDTA solution after 1 hour of encapsulation on Day 1, and another set (n=3, mean as column height) of PEARLs were lysed and recovered using 25 mM EDTA solution after 14 h days of encapsulation on Day 15 **c)** Plot showing recovery of bacteria (wcfs1 $P_{tec}$_3050 nuc) from PEARLs incubated in ZF medium for 14 days (n=3, mean as column height); on the $15^{th}$ day, PEARLs were incubated in 25 mM EDTA solution for 30 minutes with shaking, followed by plating of spun down bacterial pellet resuspended in MRS nutrient broth; **d)** Images of DNase + methyl green agar plates with bacteria streaked from randomly selected colonies recovered from PEARLs; Halo formation for most of the recovered colonies indicates the active secretion of nuclease.
